# SMARCA4 regulates spatially restricted metabolic plasticity in 3D multicellular tissue

**DOI:** 10.1101/2021.03.21.436346

**Authors:** Katerina Cermakova, Eric A. Smith, Yuen San Chan, Mario Loeza Cabrera, Courtney Chambers, Maria I. Jarvis, Lukas M. Simon, Yuan Xu, Abhinav Jain, Nagireddy Putluri, Rui Chen, R. Taylor Ripley, Omid Veiseh, H. Courtney Hodges

**Affiliations:** Department of Molecular and Cellular Biology, Baylor College of Medicine, Houston, TX, USA; Center for Precision Environmental Health, Baylor College of Medicine, Houston, TX, USA; Department of Bioengineering, Rice University, Houston, TX, USA; Therapeutic Innovation Center, Baylor College of Medicine, Houston, TX, USA; Department of Surgery, Division of General Thoracic Surgery, Baylor College of Medicine, Houston, TX, USA; Department of Epigenetics and Molecular Carcinogenesis, The University of Texas MD Anderson Cancer Center, Houston, TX, USA; Department of Molecular and Human Genetics, Baylor College of Medicine, Houston, TX, USA; Center for Cancer Epigenetics, The University of Texas MD Anderson Cancer Center, Houston, TX, USA; Dan L Duncan Comprehensive Cancer Center, Baylor College of Medicine, Houston, TX, USA

**Author notes:** These authors contributed equally to this work (Katerina Cermakova, Eric A. Smith). Corresponding author: H. Courtney Hodges, Department of Molecular and Cellular Biology, Baylor College of Medicine, One Baylor Plaza, Houston, TX, USA,.

## Abstract

SWI/SNF and related chromatin remodeling complexes act as tissue-specific tumor suppressors and are frequently inactivated in different cancers. Although many regulatory activities of SWI/SNF have been identified using 2D cell culture, the effects of SWI/SNF alterations in more complex 3D tissues have remained poorly understood. Here we employed 3D cell culture conditions that yield transcriptomic states mirroring primary lung adenocarcinoma (LUAD) specimens better than 2D culture. By analyzing spatial patterns of gene expression and DNA accessibility in 3D spheroids using single-cell RNA-seq and ATAC-seq, we find that the SWI/SNF ATPase SMARCA4 (BRG1) induces state-specific changes to DNA accessibility that influence spatially heterogeneous expression patterns and metabolism. In 3D conditions, SMARCA4 promotes accessibility for AP-1 transcription factors, including ATF3, a regulator of metabolism and repressor of NRF2 antioxidant signaling. These changes reduce expression of SLC7A11 in a distinct portion of cells, which sensitizes A549 spheroids to cell death via ferroptosis under oxidizing conditions. Consistent with these results, we find that SMARCA4 alterations are associated with derepression of NRF2 targets in human tumors independently of NRF2/KEAP1 status. Our work reveals new 3D-specific features and unanticipated spatial complexity associated with chromatin remodeling in multicellular tissues.

## Introduction

A large number of cellular signals are processed via integration on the chromatin landscape^1,2^. As a result, chromatin-based mechanisms are essential to normal cell function, and alterations of these mechanisms have major contributions to diverse human diseases. SWI/SNF family ATP-dependent chromatin remodelers, which use ATP to physically remodel histones and other factors on chromatin^3^, generate DNA accessibility for a wide variety of transcription factors^4–6^ and other DNA-templated processes. In recent years, sequencing of genetic and epigenetic alterations associated with disease states has revealed that SWI/SNF subunits are among the most frequently disrupted genes in a number of diseases^7–9^, including neurodevelopmental disorders^10,11^, autism spectrum disorders^12,13^, and cancer^14^.

SWI/SNF complexes enable transcriptional activity that varies in time, a key feature of cellular plasticity. Temporally variable activities regulated by SWI/SNF are essential for rapid response to many intracellular and extracellular signals, including differentiation^15,16^, immune activation^17^, cell cycle exit^18^, and maintenance of metabolic homeostasis^19,20^. Consistent with its contributions towards rapid chromatin state change, chemical-induced recruitment of SWI/SNF complexes to a chromatin locus results in local eviction of repressive factors within minutes^21^. Moreover, loss of SWI/SNF activity induces immediate alterations of the chromatin landscape, indicating that remodeling activity is employed in an ongoing manner to generate DNA accessibility^22,23^ and plays an important role in maintenance of steady-state chromatin profiles.

In contrast to temporal features, the spatial variability arising from chromatin remodeling activity in more complex 3D tissue has remained less well understood. SWI/SNF and related complexes are altered in ~20% of all cancers^7^, and frequently act as tissue-specific tumor suppressors^14,24^. Unlike 2D cell culture, most tumors are spatially heterogenous 3D tissues that exhibit a high degree of phenotypic zoning^25,26^. Such spatially variable features include regions of differential proliferation, hypoxia, and/or nutrient utilization^27–29^. Unfortunately, the interplay between SWI/SNF activity and variable 3D conditions has remained uncertain. Improved experimental models that mimic 3D-specific metabolic conditions^25,26^ while limiting cell-type heterogeneity would provide a new opportunity to investigate the activities of SWI/SNF in an experimentally controlled 3D tissue.

Here we have employed 3D cell culture, which permits the study of SMARCA4’s effects on chromatin and transcription across a fuller spectrum of conditions expected to arise in vivo. Unlike animal models, which show a high degree of heterogeneity between and within tumors, 3D spheroid models give rise to heterogeneous spatial features while providing a higher degree of experimental control^30^. We employed lung adenocarcinoma (LUAD) spheroids, because lung cancers are characterized by a high degree of non-genetic heterogeneity^31,32^. LUAD cells represent especially convenient 3D tissue model, as they spontaneously form 3D multi-cellular tumor spheroids (MCTSs or “spheroids”) in non-adherent conditions, which establish spatially variable states via differential exposure of tumor cells to oxygen and nutrients. In LUAD, SWI/SNF subunits are mutated in ~17% of cases^33^, shared between the ATPase SMARCA4 (BRG1) and ARID1A. The activity of SMARCA4 in LUAD has furthermore been linked to regulation of oxidative phosphorylation and to repression of the antioxidant NRF2 pathway^34^, however chromatin mechanisms by which SMARCA4 contributes to oncogenesis and tumor development have remained unresolved.

By examining the spatial dependence of SMARCA4-induced gene expression and DNA accessibility, we have uncovered that SMARCA4 promotes DNA accessibility in a manner that is responsive to metabolic adaptations in distinct portions of 3D tissue. We find that SMARCA4 activity is surprisingly not static or invariant, but occurs in a state-dependent manner that is strongly influenced by nutrient availability and metabolic conditions. Our results show that SMARCA4 activity constrains metabolic plasticity by promoting accessibility for transcription factors involved in metabolism and stress response, and suggest that SMARCA4 alterations permit LUAD cells to mount a fuller response to oxidative stress conditions.

## Results

### Three-dimensional spheroid cultures are more faithful models of human LUAD tumors

Cells in 2D monolayers and 3D tissues display distinct dependencies^35^. To test whether MCTSs better resemble genome-wide expression patterns of human LUAD tumors than cells cultured in 2D, we prepared spheroids from seven LUAD cell lines: A549, H1299, H2030, H1975, H23, H358, and H441 (**Figures 1a**, details in Methods). All seven cell lines spontaneously formed spherical multicellular tissues in non-adherent conditions (**Figure 1b**). The high self-adherence of these cell lines is common amongst cells of the lung^36^, and consistent with primary LUAD tumors, which often show large nodules of dense transformed cells without strong intermixing with stroma^37^.

**Figure 1.**
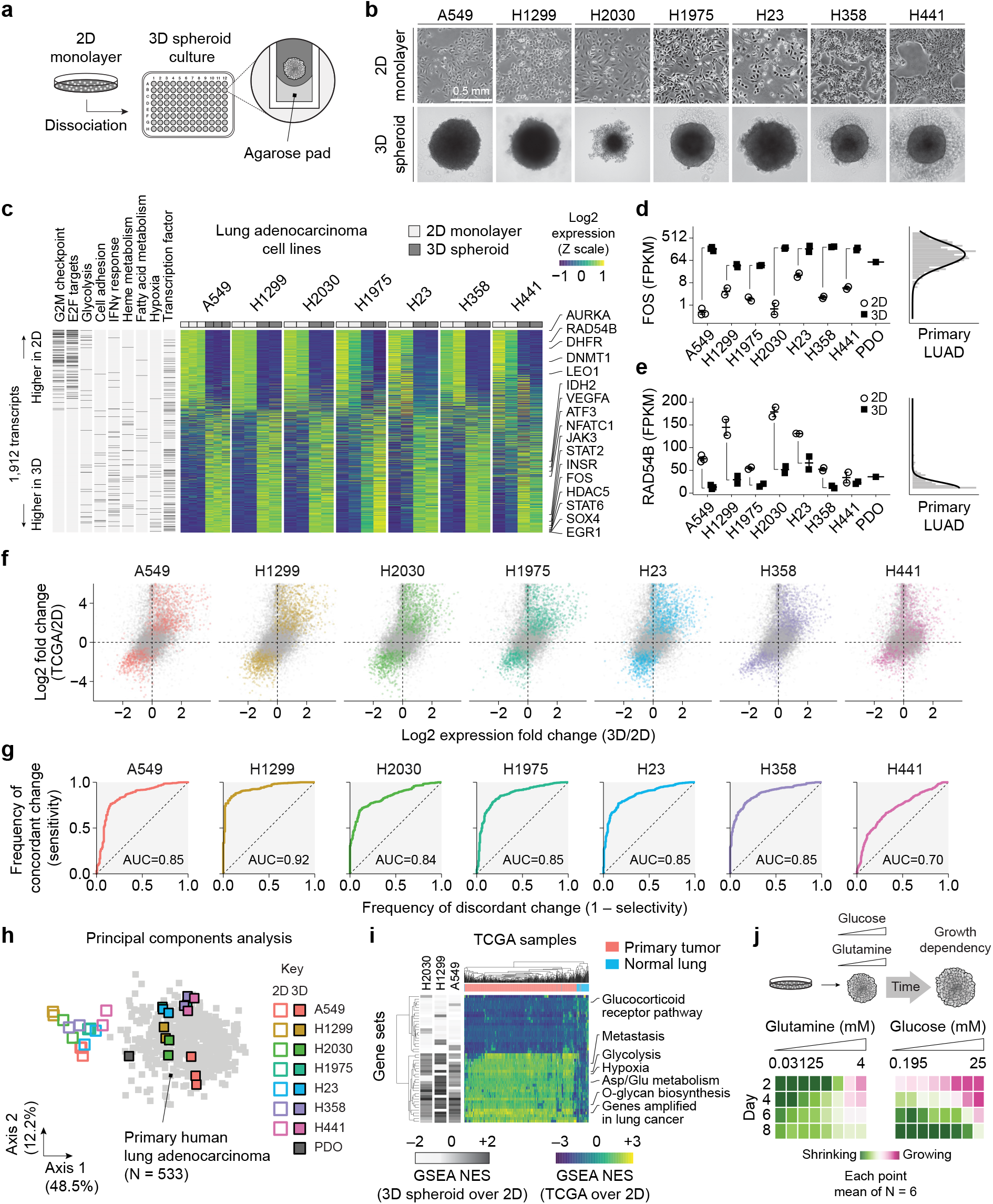
3D LUAD spheroids recapitulate transcriptomic and metabolic features of primary tumors. **(a)** Assembly of 3D spheroids. **(b)** Brightfield images of 2D monolayer and 3D spheroids derived from seven LUAD cell lines. **(c)** Signature of 1,912 genes with 3D-specific expression patterns in seven LUAD cell lines. **(d)** Expression of FOS in 2D and 3D cultured LUAD cell lines compared to human tumors. **(e)** Expression of RAD54B in 2D and 3D cultured LUAD cell lines compared to human tumors. **(f)** Expression changes induced by 3D culture compared to changes between 2D and TCGA specimens. Genes highlighted in colored points are members of the signature identified in panel (c). **(g)** ROC curves assessing expression patterns induced by 3D culture condition with primary tumors. Area under the curve (AUC) metric for each curve is presented. **(h)** Principal components analysis of transcriptomes of LUAD cell lines cultured in 2D vs. 3D, overlaid with PDO and TCGA LUAD patient tumors. **(i)** Heatmap of GSEA scores for 3D vs. 2D LUAD cell line transcriptomes compared to TCGA LUAD patient tumor and normal lung tissue transcriptomes. See also **Figure S1**. **(j)** Change in A549 spheroid growth over time in varying concentrations of glutamine and glucose. Abbreviations: FPKM, fragments per kilobase of transcript per million mapped reads; ROC, receiver operating characteristic; PDO, patient-derived organoid.

To examine genome-wide transcriptional differences between 2D and 3D cultured LUAD cell lines, we performed RNA-seq of all seven cell lines in both conditions. All spheroid models retained distinguishing features, but also adopted 3D-specific expression patterns (**Figure S1a-c**). Within each individual line, the changes induced by 3D culture showed a high degree of correspondence to the differences between 2D cells and primary human tumors from The Cancer Genome Atlas (TCGA, see example in **Figure S1b**). As we observed correlated expression changes in all 3D models, we sought to identify whether a common 3D-specific signature was shared by all lines. Using a fault-tolerant strategy to identify overlapping genes (described in Methods), we uncovered a signature of 1,912 transcripts with recurrent changes between 2D and 3D conditions (**Figure 1c** and **Supplemental Dataset 1**). This signature spanned a broad range of functions and included genes involved in proliferation, metabolism, and transcription factors. Several genes, such as the AP-1 transcription factor FOS or RAD54B, were up- (**Figure 1d**) or down-regulated (**Figure 1e**) to expression levels in quantitative agreement with primary tumors.

3D-induced expression changes mirrored the principal differences between 2D cells and primary tumors (**Figure 1f**, Pearson’s product-moment correlation test, P<2.2e-16 for all seven cell types). To assess the proportion of concordant and discordant changes between 3D conditions and primary tumors for each line, we plotted receiver operating characteristic (ROC) curves and found that expression changes between 2D and 3D states were predictive of expression changes between 2D and primary tumors with high sensitivity and selectivity (**Figure 1g**). 3D samples furthermore clustered with primary samples using unbiased approaches. Principal components analysis (PCA) of all experimental samples based on all differentially expressed genes with (FDR)-adjusted P<1e-4 revealed a clear separation between 2D and 3D samples. Overlaying the transcriptomes from 533 primary TCGA LUAD tumors and a 3D patient-derived organoid (PDO) on the same axes revealed that these samples readily clustered with 3D rather than 2D samples (**Figure 1h**).

To confirm that transcriptomic changes induced by 3D culture were specific to cancer, and to rule out that these changes were a general byproduct of all cells in 3D, we analyzed differential enrichment of gene sets in 2D and 3D from A549, H1299, and H2030 cells, and measured the enrichment in primary tumors and the limited number of matched normal tissue samples from TCGA (**Figure 1i**). Gene set enrichment analysis (GSEA) revealed that gene sets upregulated in 3D spheroids are also upregulated in primary tumors and not enriched in normal tissue (**Figure 1i**). Many of gene sets enriched in 3D are involved in metabolic features (**Figure S1e-f** and **Supplemental Dataset 2**).

LUAD cells in 2D culture display strong glutamine utilization but primarily metabolize glucose in vivo^38^. To reveal whether LUAD spheroids display metabolic properties more consistent with 2D cell culture or in vivo tumors, we performed metabolic dropout studies. By monitoring spheroid growth in varying concentrations of glucose and glutamine, we found that spheroids undergo a metabolic switch between days 3-6 from being primarily dependent on glutamine, to being primarily dependent on glucose (**Figure 1j**). Mature LUAD spheroids therefore display glutamine and glucose utilization patterns that mimic in-vivo conditions and are distinct from 2D patterns. Together, our results indicate that 3D spheroid culture provides a substantially improved model of LUAD tumor cell physiology compared to conventional 2D culture.

### SMARCA4 regulates the spatial expression of genes involved in key metabolic pathways

We reasoned that improved 3D cell culture models provided a new opportunity to uncover the mechanisms by which SMARCA4 alterations in LUAD contribute to disease progression (**Figure 2a**). We used lentiviral transduction to express either SMARCA4 at physiological levels or empty vector (EV) controls in SMARCA4-null LUAD cell lines^39,40^ (**Figure 2b**), which we used to generate 3D spheroids (**Figure 2c**). Compared to EV controls, SMARCA4-expressing cells showed no discernable morphological change in 2D culture (**Figure S2a**) but displayed pronounced morphological changes in 3D spheroids as measured by high-content microscopy (**Figure S2b**). In contrast to the EV control, SMARCA4-expressing spheroids were smaller and smoother in appearance when seeded with same number of cells (**Figure 2d-e**). Hence SMARCA4 reliably induces quantifiable morphological phenotypes under 3D conditions.

**Figure 2.**
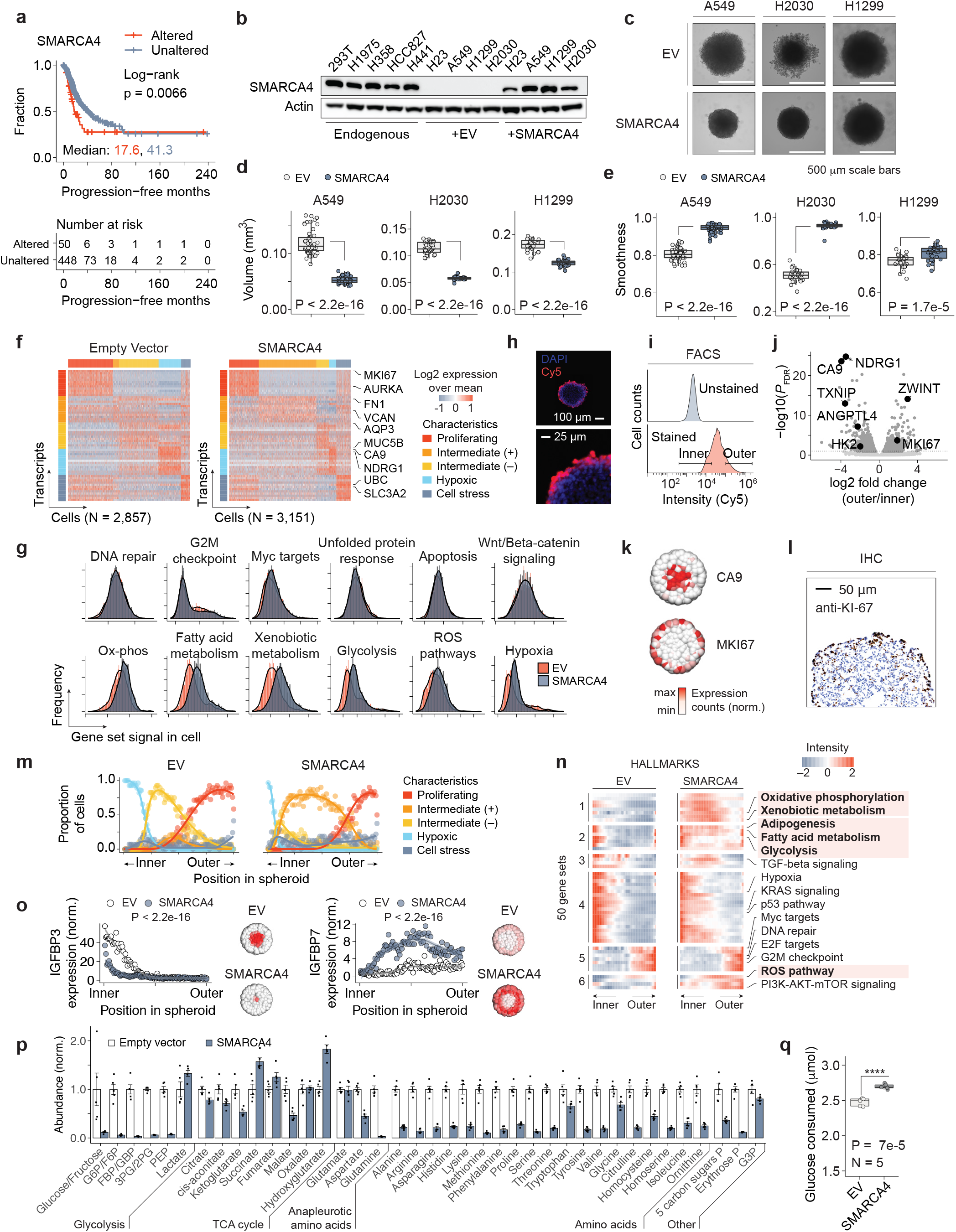
SMARCA4 regulates the spatial expression patterns of genes in key metabolic pathways. **(a)** Progression-free survival of LUAD patients based on SMARCA4 alteration status. **(b)** SMARCA4 western blot in wild-type cell lines alongside SMARCA4-null cell lines transduced with EV or SMARCA4. **(c)** Brightfield images of 3D spheroids seeded with identical number of cells expressing EV or SMARCA4. **(d)** Spheroid volume measurements of LUAD spheroids. **(e)** Edge smoothness measurements of LUAD spheroids. **(f)** Single-cell transcriptional profiles clustered into groups by K means clustering. **(g)** Single-cell enrichment histograms of cancer hallmark gene sets in SMARCA4 and EV conditions. **(h)** Fluorescence microscopy of A549 spheroid cross section after staining intact spheroid with fluorescent ester, counterstained with DAPI. **(i)** Flow cytometry of cells from unstained and stained spheroids, and gating strategy for inner and outer fractions. **(j)** Differential genes expressed in outer vs. inner spheroid fractions determined by RNA-seq. **(k)** Inferred spatial expression patterns of CA9 and KI-67. **(l)** KI-67 IHC staining of spheroid cross section to validate inferred spatial expression. **(m)** Frequency of cells from different clusters from panel (f) based on their inferred position. **(n)** Spatially resolved HALLMARK gene set enrichment for A549 spheroids. Identities of all gene sets are provided in **Supplemental Dataset 7**. **(o)** Spatial expression of individual genes in A549 spheroids and inferred expression patterns. **(p)** Steady-state metabolomic profiling in A549 spheroids. N=5, each metabolite. **(q)** Glucose consumption of A549 spheroids. For all panels: EV, Empty vector

We performed single-cell RNA-seq (scRNA-seq) on genetically monoclonal A549 spheroids (**Figure S2c-d**) to focus on SMARCA4-dependent regulation of phenotypic states rather than genetic heterogeneity. scRNA-seq revealed heterogeneous expression patterns in 3D spheroids and permitted assignment into 5 bins based on K-means clustering (**Figure 2f**). Each cluster displayed distinct characteristics, such as signatures of proliferation or hypoxia. We therefore sought to identify specific features or pathways regulated by SMARCA4 at the single-cell level, and analyzed gene sets contained within cancer hallmarks records of MSigDB^41^.

In other contexts, SWI/SNF has been observed to regulate DNA repair^42^, cell cycle progression^43–46^, Myc targets^47^, Wnt/β-catenin signaling^48^, unfolded protein response^49^, apoptosis^50^, or other classic modes of tumor suppression. However, our results at this stage did not support a prominent role for SMARCA4 in the regulation of these features in 3D LUAD cells. Instead, we observed pronounced SMARCA4-associated activation of key metabolic pathways, including oxidative phosphorylation (ox-phos), fatty acid metabolism, glycolysis, reactive oxygen species (ROS), and hypoxia (**Figure 2g**). Increased gene expression does not necessarily indicate increased metabolic activity of each pathway, as increased transcription could reflect heightened capacity, or compensation response to reduced activity. A more detailed analysis suggests that SMARCA4 has a modest effect on the proportion of G1 and G2 cells, but altered proliferation rates were not detected during the time of spheroid culture (**Figure S3d-g**). Therefore, our results indicate that in 3D LUAD spheroids, SMARCA4 has stronger gene expression effects on key metabolic pathways than traditional modes of tumor suppression.

To add spatial resolution to the scRNA-seq data, we developed a FACS-based separation approach that enabled us to identify transcriptomic features based on position/perfusion within spheroids (**Figures 2h-i** and **S2e**). We stained intact spheroids with CellTrace Far Red, a cell-permeable fluorescent ester, then dissociated cells and used FACS to separate bright fluorescent (outer) cells from dim (inner) cells (**Figures 2i** and **S2f-j**). We then obtained bulk RNA-seq signatures of each portion of the spheroids, which revealed that outer cells expressed higher levels of proliferative markers (e.g., *KI-67*), while inner cells expressed higher levels of hypoxia genes (e.g., *HK2* and *CA9*, **Figures 2j**, **S3a-b**, and **Supplemental Dataset 3**). We used standard approaches to infer the spatial position of cells within spheroids using a pseudospace analytical metric based on inner and outer markers (**Figure S3h-i**). To reduce noise, we pooled cells within 100 consecutive 1%ile bins based on their inferred positions. Our results showed that hypoxic markers (e.g. *CA9*) were spatially constrained to the inner portion of the spheroid, while proliferative markers (e.g., *KI-67*) were restricted to outer regions (**Figure 2k**). We also confirmed the differential increase of KI-67 in outer cells by IHC staining of intact spheroids (**Figure 2l**). Assigning spatial position to each cell revealed that the five clusters we previously observed at the single-cell level were not random but were spatially restricted (**Figure 2m**) and that SMARCA4-regulated gene sets were especially influenced in the intermediate zone between outer normoxic proliferating cells, and hypoxic cells on the interior of the spheroid. Spatial expression profiles for 22,115 genes in EV and SMARCA4 spheroids is provided in **Supplemental Datasets 4-5**.

In agreement, analysis of gene sets revealed pronounced spatial segregation of cancer hallmarks, including hypoxia on the interior and proliferation on the outer portions (**Figure 2n**). Our results confirm that nutrient and oxygen gradients induce spatial patterning of gene expression in 3D tissues over length scales as small as 0.5 mm and provide a new approach to analyze these gradients at the transcriptomic level. Additionally, the SMARCA4-dependent induction of key metabolic gene sets similarly showed a spatial component, with several sets (highlighted in **Figure 2n**) that were restricted to hypoxic regions in EV control cells expanding to the intermediate zones in SMARCA4-expressing spheroids. Similar results were obtained by analyzing the 186 gene sets in the KEGG database (**Figure S3k**). Importantly, all regions of the spheroid displayed SMARCA4-dependent effects, including genes in the inner and outer regions (**Figure 2o**). Our results show that the heterogeneous effects of SMARCA4 on gene expression are responsive to the spatially variable patterns of nutrient and oxygen availability.

Steady-state LC/MS-MS metabolomic profiling confirmed changes to a large number of altered metabolic intermediates of glycolysis, amino acid metabolism, and the pentose phosphate pathway (**Figure 2p**). We also confirmed differential glucose utilization by direct measurement of residual glucose in conditioned media (**Figure 2q**). Together our results show that SMARCA4 activity influences diverse metabolic pathways and is state-dependent.

### Regulation of DNA accessibility in 3D tissues by SMARCA4

To identify sites where SMARCA4 directly regulates chromatin, we developed a strategy to map the spatial patterns of DNA accessibility promoted by SMARCA4 in 3D space (**Figure 3a**). We employed a similar FACS procedure to identify ATAC-seq peaks as markers of inner and outer cells for single-cell ATAC-seq (scATAC-seq, **Figure S4a**), and separately, employed bulk ATAC-seq and ChIP-seq in SMARCA4 and EV cells as a conservative measure to identify sites regulated by SMARCA4 throughout the body of the spheroid. We limited our analysis to sites with increased DNA accessibility in SMARCA4 cells via bulk ATAC-seq (N=684) that overlapped with ChIP-seq peaks for SMARCA4 (N=20,530, **Figure S4b-c**). Overlap of these sites yielded 209 sites, reflecting significant overlap between increased sites and SMARCA4 peaks compared to all accessible sites (N=107,746, odds ratio OR=2.54, P=1.71e-13). Example browser tracks and scATAC spatial profiles are provided in **Figure 3b-c**. Spatial positions in the spheroid where these SMARCA4 target sites displayed increased accessibility were frequently constrained (**Figures 3d**, **S4d-f**), indicating that SMARCA4-dependent DNA accessibility is strongly influenced by the local cell environment, consistent with state-dependent chromatin remodeling.

**Figure 3.**
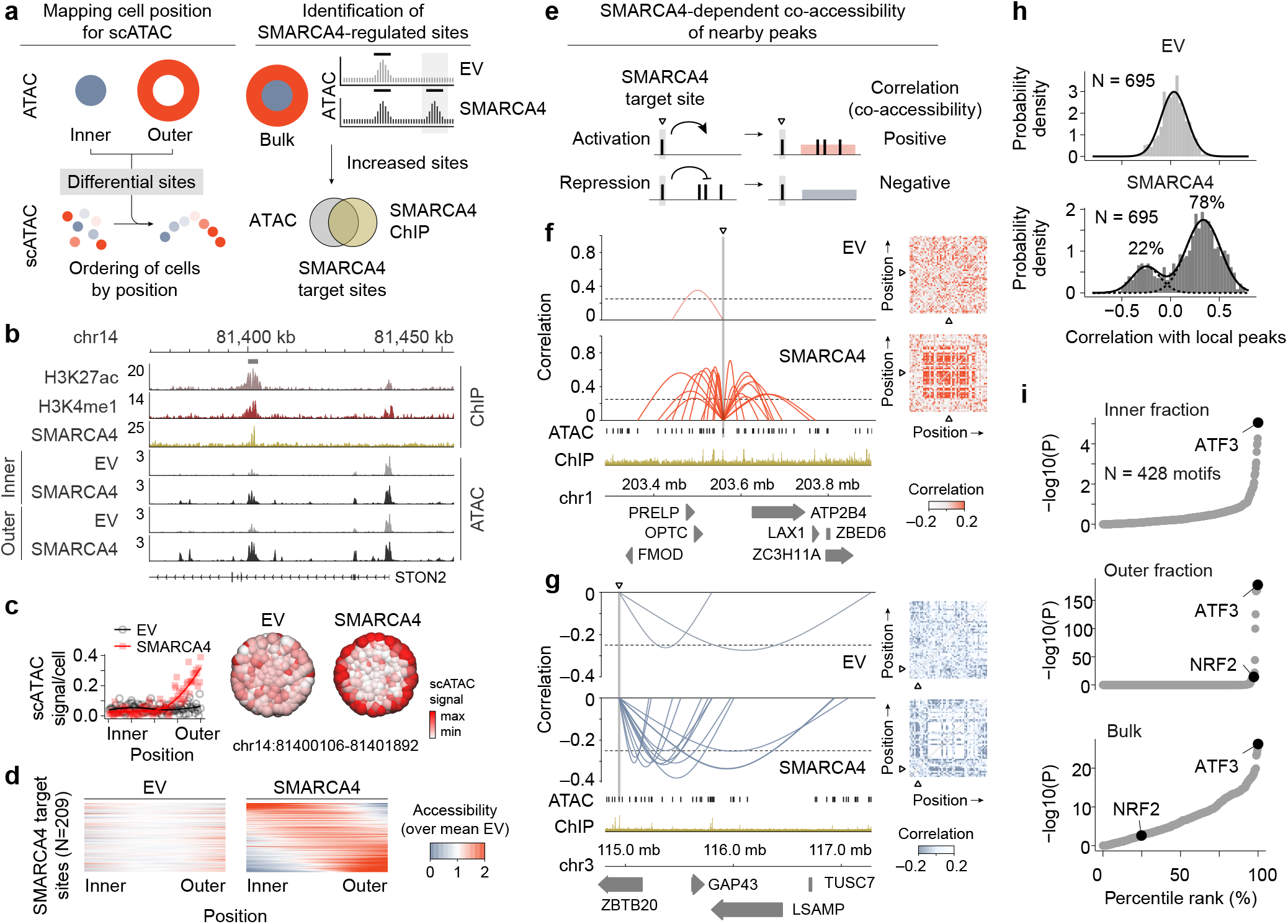
SMARCA4 regulates accessibility for ATF3 in a state-dependent manner. **(a)** Workflow for mapping of cell positions in scATAC-seq and for identification of SMARCA4 targets. **(b)** Example browser tracks comparing ATAC-seq peaks of inner and outer fraction of A549 cells expressing EV or SMARCA4 alongside H3K4me1, H3K27ac and SMARCA4 ChIP-seq profiles. **(c)** Spatially regulated accessibility profile and inferred spatial distribution regulated by SMARCA4. **(d)** Spatial distribution of accessibility for SMARCA4 target sites. **(e)** Identification of activating or repressive SMARCA4 sites within CCANs. **(f)** Example of SMARCA4-activated sites within CCAN. **(g)** Example of SMARCA4-repressed sites within CCAN. **(h)** Distribution of pairwise correlation between SMARCA4-regulated sites and other sites within CCAN boundaries. **(i)** Transcription factor motif enrichment in inner fraction, outer fraction and bulk ATAC-seq ranked by P value. For all panels: EV, Empty vector; CCAN, *cis*-co-accessibility network

To assess the impact of SMARCA4 targets on neighboring sites in the genome, we performed an analysis of correlation of SMARCA4-dependent accessibility (or co-accessibility). We identified regions of high local correlated accessibility (termed *cis*-co-accessibility networks or CCANs^51^) based on scATAC-seq of SMARCA4-expressing spheroids and investigated whether sites within these regions displayed signs of SMARCA4-dependent activation or repression (see **Figure 3e**, details in Methods). Interestingly, SMARCA4 target sites within CCANs displayed either positive correlation with other sites in these regions, consistent with activation (**Figure 3f**), or anti-correlation with other sites consistent with repression (**Figure 3g**). The pattern of locally correlated domains supports the view that SMARCA4 generates accessibility for factors that regulate local chromatin domains in *trans* through long-range contacts.

To assess the relative contributions of activation versus repression, we analyzed pairwise correlation for sites within CCANs with the respective SMARCA4 target site contained within the CCAN (N=695, **Figure 3h**). At these sites, pairwise correlation was normally distributed around zero for EV cells (Shapiro-Wilk normality test P>0.05, mean R=0.035±0.13 s.d.), but SMARCA4 cells displayed a bimodal distribution of both positive and negative values (Shapiro-Wilk normality test P=7.6e-15). The SMARCA4 distribution was readily fit by maximum-likelihood estimates of two normal distributions with mean R_1_=0.34±0.18 s.d. and R_2_=-0.25±0.14 s.d. This indicates that SMARCA4 generates accessibility for factors that activate or repress neighboring sites in *trans*. Despite a clear bias in favor of activation, ~22% of pairwise interactions displayed negative correlation, consistent with SMARCA4-dependent loading of a repressor.

We examined which transcription factor motifs are enriched in SMARCA4-increased ATAC-seq peaks compared to unchanged sites. The most frequent overall enriched motif in the inner, outer, and bulk ATAC libraries of A549 3D spheroids was the motif for binding of the AP-1 family transcription factor ATF3 (**Figure 3i**). ATF3 is a known regulator of metabolic homeostasis, and plays important roles in tumor metabolism^52^ and stress response^53,54^. ATF3 is moreover known to directly bind SMARCA4^55^, however a SMARCA4-dependent role for regulation of ATF3 in LUAD has not previously been reported. The motif for NRF2 was also significantly enriched in bulk (Hypergeometric test P=2.8e-3) and outer fraction (P=2.3e-14) of A549 spheroids but less significantly than the ATF3 motif, and unlike ATF3 was not enriched in the inner fraction (P>0.05). ATAC-seq of H2030 3D spheroids yielded comparable results (**Figure S4g**).

### SMARCA4 promotes ATF3 placement at enhancers of NRF2 target genes

Loss of SMARCA4 was recently shown to influence the expression of NRF2 targets^34^ and ATF3 is a negative regulator of NRF2 signaling^56,57^. To understand the interplay between SMARCA4, ATF3, and NRF2, we performed ChIP-seq for these factors in bulk 3D spheroids. In SMARCA4-expressing cells, we observed 2,969 ATF3 and 59 NRF2 sites (**Figure 4a**). SMARCA4 peaks overlapped with a majority of both NRF2 and ATF3 sites (**Figure 4b-c**), however few ATF3 and NRF2 sites showed overlap, consistent with their distinct binding motifs. SMARCA4 expression induced reproducible redistribution of ATF3 occupancy genome-wide (**Figure 4d-e**, Pearson Correlation test P<2.2e-16, R=0.404 across independent replicates). In particular, we observed SMARCA4-dependent increase of ATF3 occupancy at the promoter of the cystine/glutamate antiporter SLC7A11 in both ChIP-seq (**Figure 4d**) and ChIP-qPCR (**Figure S4h**).

**Figure 4.**
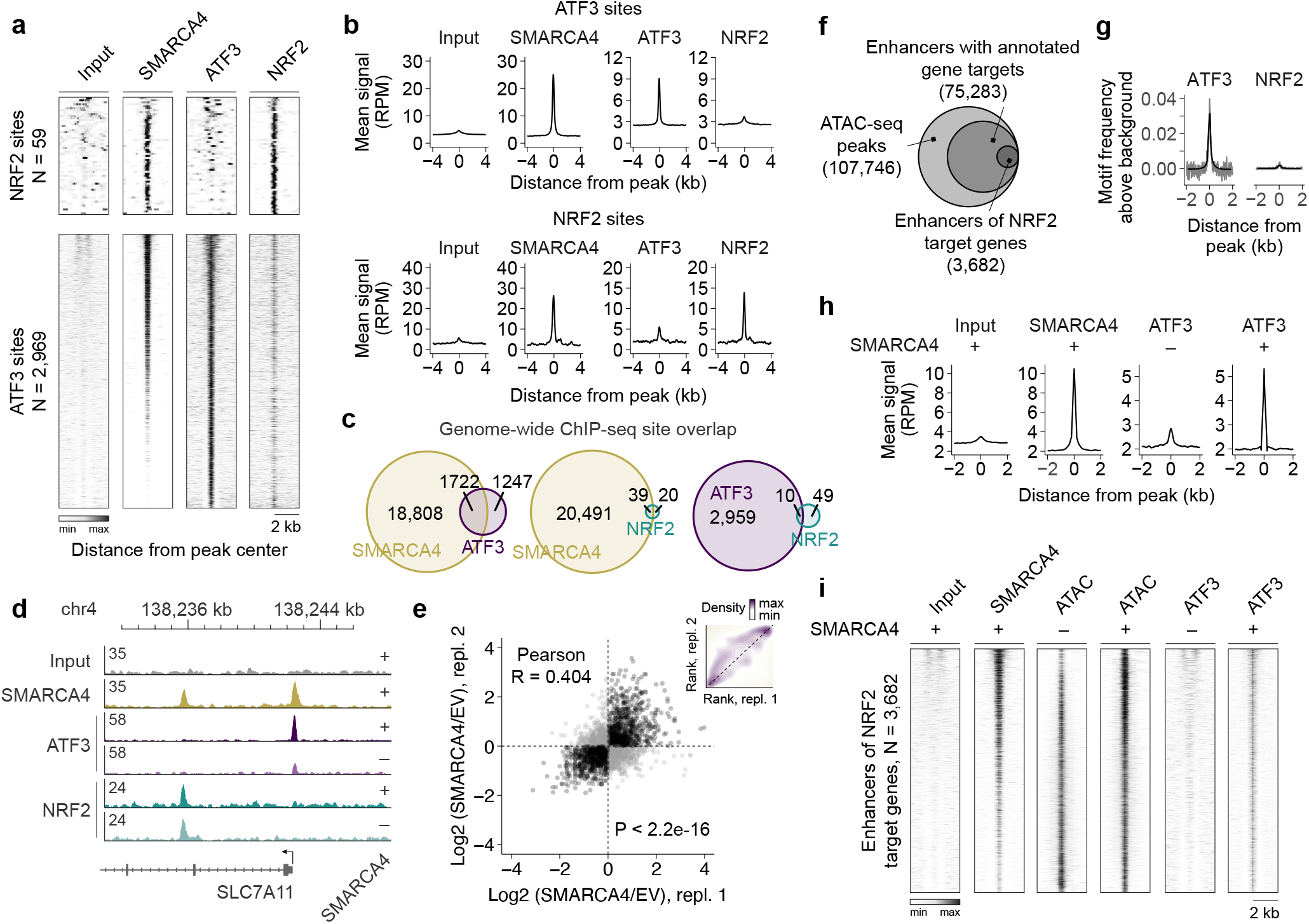
SMARCA4 promotes ATF3 binding at NRF2 target genes. **(a)** ChIP-seq of SMARCA4, ATF3, and NRF2 in bulk 3D A549 spheroids expressing SMARCA4. **(b)** Examples of ChIP peaks at SMARCA4-bound NRF2 and ATF3 sites. **(c)** Venn diagrams showing SMARCA4, NRF2 and ATF3 ChIP-seq peak overlap. **(d)** Browser tracks comparing SMARCA4, ATF3, and NRF2 ChIP-seq peaks in EV and SMARCA4 expressing A549 spheroids at the SLC7A11 locus. **(e)** Correlation of ATF3 ChIP-seq changes induced by SMARCA4 across replicates. **(f)** Identification of enhancers of NRF2 target genes. **(g)** Frequency of NRF2 and ATF3 motifs at enhancers of NRF2 target genes. **(h)** Mean intensity of SMARCA4-dependent ATF3 binding at enhancers of NRF2 target genes. **(i)** SMARCA4-dependent ATF3 binding at enhancers of NRF2 target genes. For all panels: EV, Empty vector

Because ATF3 has been shown to repress NRF2 target genes^56^, we sought to establish whether SMARCA4 promoted the placement of ATF3 at enhancers of NRF2 target genes. We identified the locations of all enhancers of NRF2 target genes by cross-referencing NRF2 targets in MSigDB with their annotated enhancers in the GeneHancer database^58^. This procedure resulted in 3,682 sites, which had higher enrichment of the ATF3 motif than NRF2 motif (**Figure 4g**) and high levels of SMARCA4 occupancy (**Figure 4h-i**). We observed a marked increase of ATF3 occupancy at these sites in SMARCA4 cells compared to EV cells (**Figure 4h-i**), thereby confirming that enhancers of NRF2 target genes are subjected to SMARCA4-dependent placement of ATF3. Because ATF3 is a known repressor of NRF2 signaling, this activity represents a novel mechanism for antagonism of NRF2 signaling at the chromatin level.

### SMARCA4 represses SLC7A11, potentiating oxidative stress and sensitivity to ferroptosis

We sought to identify and validate phenotypic consequences associated with SMARCA4-dependent chromatin regulation that may represent novel tumor suppressive activities. Ranking of SMARCA4-bound genes associated with NRF2 and ATF3 sites revealed SLC7A11 (**Figure 4d**) to be a top regulatory target for both ATF3 (98.9%ile) and NRF2 (99.0%ile, **Figure 5a**). High levels of SLC7A11 are associated with worse progression-free survival in human LUAD patients (**Figure S5a**), making it a potential vulnerability. Moreover, we observed that SLC7A11 was repressed upon SMARCA4 expression in bulk RNA-seq (**Figure 5b, Supplemental Datasets 6**), as well as in our single-cell data (**Figure S5b-e**) of 3D spheroids derived from A549 cells. We confirmed SMARCA4-dependent repression of SLC7A11 at the RNA level via RT-qPCR in 3D spheroids derived from A549, H1299, and H2030 cells (**Figure 5c**). These changes were 3D-specific, as they were not observed in 2D monolayer culture (**Figure S5f**), consistent with the low levels of ATF3 in 2D cells (**Figure 1c**). RNA levels furthermore coincided with SMARCA4-dependent reduction of SLC7A11 at the protein level as measured by Western blot in spheroids derived from all three cell lines (**Figures 5d-e** and **S5g**).

**Figure 5.**
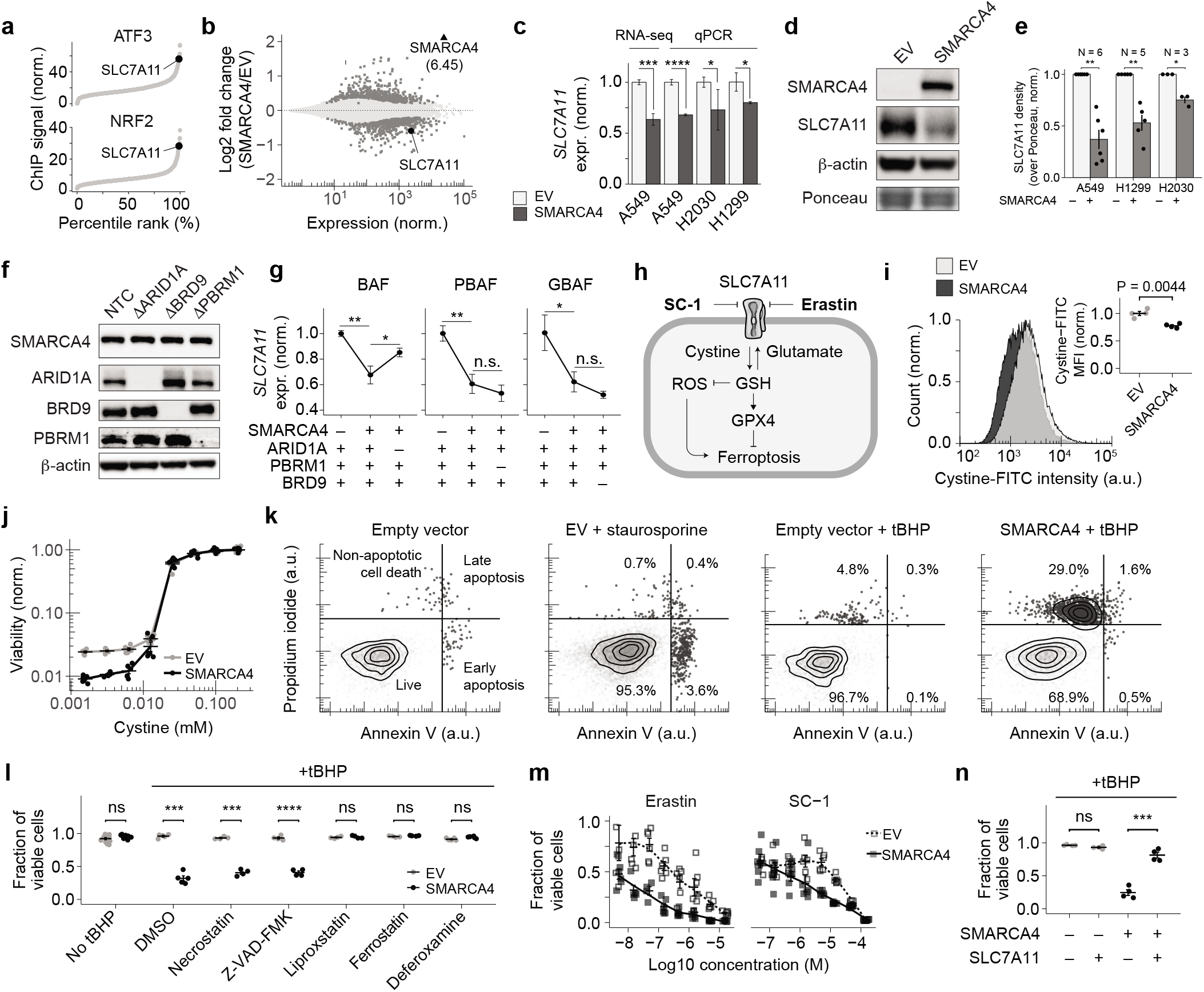
SMARCA4 regulates SLC7A11 and increases sensitivity to ferroptosis in LUAD spheroids. **(a)** Rank of gene targets of ATF3 and NRF2 by ChIP peak intensity. **(b)** SMARCA4-dependent RNA-seq expression changes for A549 spheroids. **(c)** qPCR validation of SMARCA4-dependent SLC7A11 regulation in LUAD spheroids. **(d)** Western blot of SLC7A11 expression in A549 spheroids. Other cell lines shown in **Figure S5g**. **(e)** Densitometry of SLC7A11 expression measured by western blot in A549, H2030 and H1299 spheroids. **(f)** Western blot validation of CRISPR/Cas9-mediated knockouts. NTC, non-targeting gRNA control. **(g)** SLC7A11 qPCR comparison of complex-specific knockouts in SMARCA4 re-expressing (“+”) A549 spheroids compared to EV (“–”). N=3. **(h)** Schematic of SLC7A11’s role in ferroptosis with small-molecule inducers of ferroptosis in bold. GSH, glutathione; ROS, reactive oxygen species **(i)** Cystine-FITC uptake measured by flow cytometry. **(j)** Effect of cystine dropout on cell viability after 9 days. **(k)** Assessment of apoptosis via flow cytometry with Annexin V and propidium iodide. **(l)** Viability and rescue of cell death in 400 μM tBHP with inhibitors measured by trypan blue. **(m)** Dose-response curves following treatment with erastin or SC-1. **(n)** Viability with or without ectopic SLC711 expression in 400 μM tBHP measured by trypan blue. For all panels: EV, Empty vector; tBHP, *tert*-butyl hydroperoxide; ****P < 0.0001; ***P < 0.001; **P < 0.01; *P < 0.05

SMARCA4 is present in three major SWI/SNF-related complexes: canonical BAF, PBAF, or GBAF^59–63^. To identify which of these complexes is responsible for the reduced expression of SLC7A11, we generated isogenic SMARCA4-expressing A549 knockout lines of ARID1A (BAF), PBRM1 (PBAF), and BRD9 (GBAF, **Figure 5f**). We confirmed that ARID1A knockout permitted increased SLC7A11 expression in SMARCA4-expressing cells, unlike knockout of PBRM1 or BRD9 (**Figure 5g**, P=0.013). We furthermore observed that loss of ARID1A in SMARCA4-expressing spheroids restored rough morphology present in SMARCA4-null spheroids (**Figures S5h-i**). Because ARID1A but not PBRM1 or BRD9 is epistatic to SMARCA4, we conclude that SMARCA4-dependent regulation of SLC7A11 arises in ARID1A-containing canonical BAF complexes. Interestingly, the only examined 3D spheroid model where SMARCA4-dependent downregulation of SLC7A11 did not occur was in H23 cells, which are SMARCA2-null and have homozygous deletions of BRD9 (**Figure S5j-m**). Together these results demonstrate that other alterations, particularly those affecting SWI/SNF subunits, can influence the SMARCA4-dependent regulation of SLC7A11.

SLC7A11 is a principal importer of cystine, the oxidized form of cysteine and precursor of glutathione, which is used as an antioxidant within cells^64^. Reduced antioxidant response permits accumulation of reactive oxygen species (ROS) that can result in lipid peroxidation. If left unchecked, excessive lipid peroxidation leads to programmed cell death known as ferroptosis^65^. Ferroptosis is agonized by the inhibitors of SLC7A11 erastin^65^ and SC-1^66^ (**Figure 5h**). Cell death via ferroptosis is also impaired by the antioxidants ferrostatin and liproxstatin^67^, as well as by the iron chelator deferoxamine^65^.

We confirmed that SMARCA4-dependent reduction of SLC7A11 levels led to reduced cystine uptake (T test P=0.0044, N=3, **Figure 5i**), impaired survival in low-cystine conditions (**Figure 5j**), and accumulation of ROS (T test P=7.2e-5, N=25 for each condition, **Figure S5p**). These observations led us to examine whether the increased ROS levels arising upon SMARCA4 expression compromised the capacity of cells to perform NAD(P)H-dependent reduction. Cells from 3D spheroids derived from A549, H2030, and H1299 cells all showed significantly compromised capacity for NAD(P)H-dependent reduction of resazurin upon SMARCA4 expression, consistent with increased oxidative stress (**Figure S5q**). Together our results demonstrate that SMARCA4-dependent repression of SLC7A11 functionally impairs SLC7A11-related activities.

Because the lung is a highly oxidizing environment^68^, we reasoned that the SMARCA4-dependent reduction of SLC7A11 may sensitize cells to cell death under oxidizing conditions. Indeed, spheroids expressing SMARCA4 but not EV, showed a rapid loss of viability upon exposure to *tert*-butyl hydroperoxide (tBHP, **Figure S5r**), an agent frequently used to mimic the oxidizing conditions found within lung tissues^69,70^. To ascertain the cell death pathway, we performed Annexin V/propidium iodide staining following tBHP treatment and discovered that SMARCA4 but not EV expression induced non-apoptotic cell death (**Figure 5k**). This effect was distinct from apoptosis induced by staurosporine.

To establish whether SMARCA4 led to increased lipid peroxidation, we exposed cells to C11-BODIPY, a fluorescent dye that is sensitive to oxidation by lipid hydroperoxides. We observed a significant reduction (T test P<0.001, N=9) of the fraction of reduced BODIPY in SMARCA4 cells, indicating increased lipid peroxidation (**Figure S5s**). We sought to rescue cell death using key inhibitors of cell death pathways, including DMSO vehicle control, necrostatin (inhibitor of necroptosis), Z-VAD-FMK (inhibitor of apoptosis), liproxstatin and ferrostatin (both antioxidant inhibitors of ferroptosis), and deferoxamine (iron chelator and inhibitor of ferroptosis). While DMSO, necrostatin, and Z-VAD-FMK all failed to rescue cell death induced by SMARCA4 under oxidizing conditions, liproxstatin, ferrostatin, and deferoxamine all showed complete rescue of viability (**Figure 5l** and **S5t**), affirming that the cell death pathway promoted by SMARCA4 is ferroptosis.

SMARCA4-expressing spheroids moreover displayed increased sensitivity to erastin and SC-1, both inhibitors of SLC7A11 (**Figure 5m**). These results suggested that the reduced levels of SLC7A11 in SMARCA4-expressing cells might be responsible for the increased sensitivity to ferroptosis. We confirmed this by ectopically maintaining high levels of SLC7A11 in EV and SMARCA4 cells using lentiviral transduction (**Figure S5u**). High levels of SLC7A11 rescued the rough phenotype of SMARCA4-expressing 3D spheroids (**Figure S5v**), as well as altered steady-state levels of metabolic intermediates of the pentose phosphate pathway (**Figure S6**). However, many other metabolites remained altered despite high levels of SLC7A11, indicating that SMARCA4 alters metabolic conditions upstream of its repression of SLC7A11. Most importantly, ectopic SLC7A11 expression caused near-complete rescue of SMARCA4-dependent ferroptosis in oxidizing conditions (**Figure 5n**). We conclude that the SMARCA4-dependent repression of SLC7A11 can contribute to cell death under oxidizing conditions, providing a novel mechanism that may contribute to tumor suppression.

### SLC7A11 is derepressed in SMARCA4-altered LUAD tumors in humans

To assess whether SMARCA4-dependent repression of SLC7A11 occurs in human tumors, we identified 457 TCGA LUAD tumor specimens with unaltered SMARCA4, and 46 patients with altered SMARCA4 (inclusion criteria defined in the Methods section). In agreement with our observations, altered tumors had ~3.8-fold higher median SLC7A11 expression than unaltered tumors (Wilcox Rank-sum test P=7.5e-7, **Figure 6a**). We separated and compared genome-wide transcriptomic differences between altered and unaltered tumors using GSEA, which revealed the signature of increased NRF2 target gene expression in SMARCA4-altered tumors (P=9.1e-6 and P=4.0e-5 for two different gene sets, **Figure 6b**), consistent with a loss of SMARCA4-mediated repression of these genes. To test whether regulatory elements of these genes had increased ATF3 levels in our 3D spheroids, we selected genes that were in the leading edge of increased genes in the NFE2L2.V2 gene set in human patients and also were the closest gene to a site with increased ATF3 occupancy in our 3D spheroids. This analysis yielded 8 ATF3/NRF2-regulated genes with significantly increased expression in SMARCA4-altered tumors, including SLC7A11 (**Figure 6c**, FDR-adjusted T test P<0.01 each), consistent with the SMARCA4/ATF3 repression mechanisms we have described.

**Figure 6.**
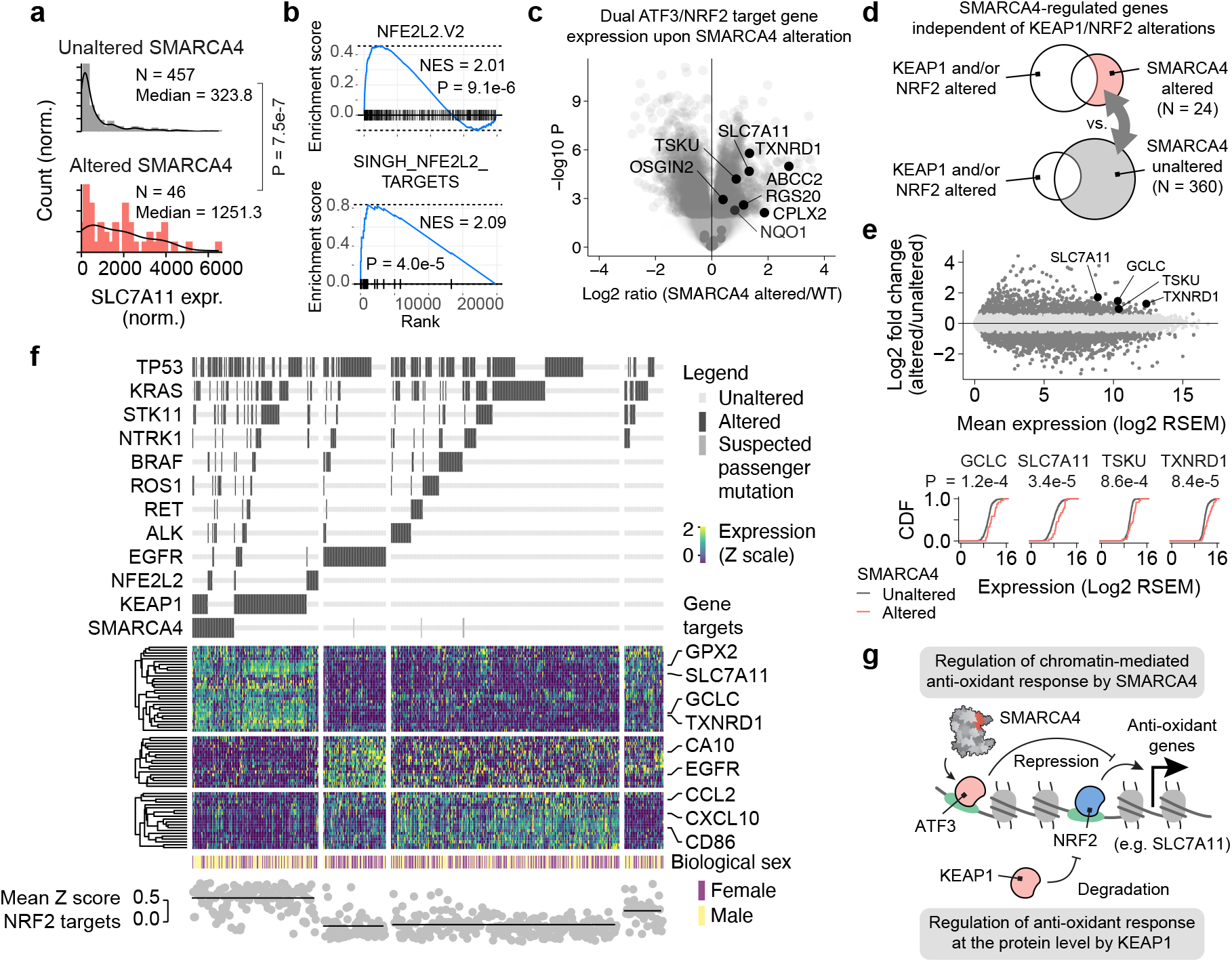
SMARCA4 alterations derepress SLC7A11 and other NRF2 targets in human LUAD tumors. **(a)** SLC7A11 expression in LUAD tumors with altered and unaltered SMARCA4. **(b)** GSEA of NRF2 targets in LUAD tumors with altered and unaltered SMARCA4. **(c)** Expression changes of ATF3-regulated genes in LUAD patients with altered and unaltered SMARCA4. **(d)** Removal of KEAP1/NRF2-mutant tumors from the cohort to investigate the SMARCA4 role in LUAD independently of KEAP1 or NRF2 mutation. **(e)** KEAP1/NRF2-wildtype RNA-seq expression changes for LUAD tumors with altered and unaltered SMARCA4. **(f)** Heatmap of selected genes highlighting mutation-specific expression features of human LUAD tumors. **(g)** Model and interpretation.

To rule out unanticipated covariates that could confound these observations, such as the presence of other mutations, we examined whether SMARCA4 alterations have the capacity to derepress SLC7A11 and other genes independently of KEAP1 alterations, which are known to activate NRF2 signaling^71^. We therefore removed KEAP1- and NRF2-mutated tumors from our cohort (**Figure 6d**) and re-examined genome-wide expression patterns. Even in KEAP1/NRF2-wild-type tumors, SMARCA4 alterations were associated with significantly increased expression of NRF2 target genes (**Figure 6e**), such as GCLC (Wilcox Rank-sum test P=1.2e-4), SLC7A11 (P=3.4e-5), TSKU (P=8.6e-4), and TXNRD1 (P=8.4e-5).

We compactly summarize our findings in a heatmap of selected genes showing mutation-specific expression features of human LUAD tumors (**Figure 6f**). In this analysis, tumors with SMARCA4 alterations show similarly elevated but visibly distinct patterns of transcriptomic features compared to tumors with KEAP1 alterations even in the absence of KEAP1 alterations (top cluster). In contrast, SMARCA4-altered tumors do not show expression features unique to EGFR alterations (middle), or inflammatory signaling (bottom). Together our analyses indicate that SMARCA4 contributes to the repression of metabolic and oxidative stress genes in human tumors even in the absence of KEAP1 or NRF2 mutations, revealing a new, 3D-specific chromatin-based mechanism for regulation of metabolic plasticity and oxidative stress.

## Discussion

There is growing recognition that 3D tumor models better mimic in vivo conditions^30,72,73^. Here we show that the transcriptomic features of primary human LUAD tumors cluster with 3D spheroids in a recurrent 3D-specific pattern, which does not arise in LUAD cell lines cultured under 2D conditions (**Figure 1**). We have also shown that 3D LUAD spheroids spontaneously establish strong patterns of non-genetic heterogeneity, owing to spatially restricted metabolic conditions (**Figure 2**). 3D spheroids therefore provide new avenues for testing hypotheses a in fuller spectrum of phenotypic states that contribute to tumors in vivo.

Although LUAD is one of the most frequent sites of SWI/SNF disruption based on incidence, a tumor suppressive role for SMARCA4 in LUAD has remained elusive. Major progress has been made on this question in recent years. The discovery that SMARCA4 influences oxidative phosphorylation in LUAD^33^ revealed that altered metabolism is a targetable consequence of SMARCA4 loss and prompted reevaluation of the links between chromatin remodeling and metabolism. Moreover, the recent observation that NRF2 signaling is altered by SMARCA4 loss^34^ provided strong evidence that constraining oxidative stress may be an important tumor suppressive activity in LUAD that contributes to cells’ metabolic strategies. However, specific chromatin regulatory mechanisms responsible for these observations have remained poorly understood.

Here we uncovered that SMARCA4 activity mediates context-dependent changes of gene expression that contribute to regulation of diverse metabolic pathways (**Figure 2**). In 3D spheroids, SMARCA4 promotes DNA accessibility for ATF3 (**Figure 3**), a factor that represses NRF2 anti-oxidant signaling^56^. The de-repression of NRF2 targets arises in SMARCA4-altered tumors from human patients independently of mutations of KEAP1, a major negative regulator of NRF2 signaling (**Figure 6**). Unlike KEAP1, which heightens antioxidant response by impairing proteolytic degradation of NRF2^74^, SMARCA4 activity opposes NRF2 activity at the chromatin level (**Figure 6g**). Our results affirm the value of 3D cell culture models, and provide a mechanistic basis for recent work showing a link between SWI/SNF activity with NRF2 signaling^34^. We note that the mechanisms we describe may be obscured or confounded by 2D cell culture, where ATF3 is typically expressed at lower levels than in 3D conditions (**Figure 1c**). We also demonstrate that SMARCA4-dependent downregulation of the NRF2 target gene SLC7A11 can be sufficient to induce cell death via ferroptosis under oxidizing conditions (**Figure 5**). Consistent with this notion, the spatial organization of NRF2 signaling was recently shown to play an important role in responding to oxidative stress^75^. Because SMARCA4 and KEAP1 are both frequently inactivated in LUAD, it is appealing to speculate that KEAP1 and SMARCA4 alterations may be under positive selection in LUAD tumors in part because their inactivation permits elevated oxidative stress response within the oxidizing environment of the lung through distinct mechanisms.

SWI/SNF-related disruptions frequently confer phenotypes that advantage tumor cells in a cell-type-specific manner. For example, ARID1A loss in ER-positive breast cancer permits tumor cells to evade estrogen deprivation therapy^76,77^, and PBRM1 loss in renal carcinoma confers resistance to immune surveillance^78^. In other cell types, alterations of SWI/SNF subunits yield different effects that promote cell survival and proliferation^79–81^. For this reason, SWI/SNF activity is often viewed to be cell-type specific. In addition to these lineage-specific functions, our results show that SWI/SNF activity is also strongly dictated by the cellular environment within a given cell type, and likely influenced by the detailed transcriptional states of each cell.

Our work reveals that SWI/SNF complexes regulate a spectrum of state-specific activities in more complex tissues, which has several important implications. Rather than a single, cell-type-specific function, the tumor suppressive role of SWI/SNF may be context-dependent, or represented by a spectrum of different effects induced by variable environmental conditions. Additionally, the positive selection for SWI/SNF inactivation may differ between tumor initiation and later stages when tumor size increases and yields hypoxic, poorly vascularized regions, or upon metastasis. Therefore, an assessment of the full spectrum of SWI/SNF activities in heterogenous tumor tissues may aid in the development of therapeutic strategies based on SWI/SNF alterations.

## Methods

### Cell lines and culture

Human cell lines were acquired from ATCC and cultured in a humidified incubator maintained at 37 °C and 5% CO2. Cell lines were maintained in high-glucose DMEM (Gibco) supplemented with 10% heat-inactivated FBS (Corning), 1X GlutaMAX (Gibco), 1X non-essential amino acids (Gibco), 1 mM sodium pyruvate (Gibco), 10 mM HEPES (Gibco), and 1X Pen/Strep (Gibco).

### Cell line generation

Lentiviruses were produced by co-transfection of Lenti-X 293T cells with packaging plasmids (psPAX2, pMD2.G) using polyethyleneimine (PEI). Following transfection, media was changed to DMEM prepared as described above but with 2% FBS. Lentiviral particles were harvested 72 hours later by passing media supernatant through a 5 μm filter (Millipore) and concentrating using a 100-kDa Amicon filter (Millipore). Cell lines were transduced using a spinfection protocol (6-well plates, 1000 x *g*, 30 minutes, 32 °C) and selected using puromycin, hygromycin, or Geneticin (G-418).

Cell lines re-expressing SMARCA4 were generated by cloning hsSMARCA4 (aka hsBRG1) from pBABE-hsBRG1 (a gift from Robert Kingston (Addgene plasmid #1959; http://n2t.net/addgene: 1959; RRID:Addgene_1959)) into a pRRL-CAG-IRES-PURO lentiviral vector. The pRRL-CAG-IRES-PURO lentiviral vector itself was used as an empty vector control. Cell lines overexpressing SLC7A11 were generated using pLenti-CMV-SLC7A11-sh926R-FLAG-IRES-Hygro (a gift from William Kaelin (Addgene plasmid #118702; http://n2t.net/addgene:118702; RRID:Addgene_118702)).Unless otherwise specified, experiments using Empty Vector and SMARCA4 re-expressing A549 cell lines refer to those lines derived from the “C9” clone of A549.

CRISPR/Cas9-mediated knockout of ARID1A, PBRM1, and BRD9 were generated using a lentiCRISPRv2 conferring Geneticin resistance (a gift from Brett Stringer (Addgene plasmid #98292; http://n2t.net/addgene:98292; RRID:Addgene_98292)). sgRNA targeting sequences are available in the supplementary table. Monoclonal isogenic cell lines harboring knockout of ARID1A, BRD9, or PBRM1 subunits in the SMARCA4 re-expressing A549 background were created by first obtaining a monoclonal SMARCA4 re-expressing A549 cell line via limiting dilution. Then, ARID1A, BRD9, or PBRM1 subunits were knocked out via lentiviral CRISPR/Cas9. Finally, monoclonal cell lines were obtained via limiting dilution and screened for knockout via Western blot.

### 3D multicellular tumor spheroid (MCTS) generation and morphometry

Multicellular tumor spheroids (MCTS) were grown in 96-well tissue culture dishes (Corning) pre-coated with 50 μL autoclaved 1% agarose (in MilliQ water). A 150 μL suspension of 10,000 cells in culture media (without any selection agent) was added to each well. Unless otherwise indicated, spheroids were cultured for 9 days prior to assay and/or collection.

Spheroid imaging was performed using the Nikon ECLIPSE Ti-2 inverted microscope for high-content imaging. Features including spheroid volume and edge smoothness were determined using the NIS Elements AR software (5.11.01) from Nikon. Morphometric parameters of live spheroids were obtained using the high-content analysis (HCA) toolbox within NIS Elements AR running on an inverted Nikon ECLIPSE Ti-2 microscope using phase contrast imaging. Spherical volume V was calculated as V = 4πr^3^/3. Smoothness S was calculated as the ratio of the length of the 2D convex hull perimeter over the length of the 2D perimeter, with both features calculated directly within NIS Elements.

### RNA preparation for bulk 2D and 3D spheroids

Total RNA was harvested from 80% confluent 2D monolayer cultures or day 9 spheroids for bulk RNA-seq. RNA for monolayer cultures was harvested using TRIzol Reagent according to manufacturer’s instructions. Spheroids were dissociated into a single-cell suspension by washing once with PBS and dissociating with TrypLE Express. RNA was subsequently harvested using TRIzol Reagent according to the manufacturer’s instructions.

### Establishment and culture of PDOs from lung adenocarcinoma cancers

Tumor samples, obtained by biopsy or surgical resection, were placed on ice in serum-free RPMI (Corning, 15-040-CV) prior to processing. Tissues were minced and dissociated using the gentleMACS system following the manufacturer’s protocol. After dissociation, the suspension was filtered through a 70 um cell strainer (BD Falcon) and washed with PBS. Dissociated cells were collected in Advanced DMEM/F12 (Thermo Fisher Scientific), pelleted, re-suspended in growth factor reduced (GFR) matrigel (Corning), and seeded in a well of a 24 well flat bottom cell culture plate (Corning). The matrigel was then solidified by a 20-minute incubation in a 37 °C and 5% CO2 cell culture incubator, and overlaid with 500 μl of DMEM/F12 media; media was subsequently refreshed every two days. The sizes and morphology was monitored daily and images of organoids with LIVE/DEAD stain were obtained after day 7 to confirm cell viability.

### Real-time quantitative PCR (RT-qPCR)

cDNA was synthesized from total RNA using the HighCapacity cDNA Synthesis Kit (Applied Biosystems), and gene expression was analyzed using the QuantStudio 3 instrument (Applied Biosystems) using TaqMan Fast Advance Master Mix (Applied Biosystems #4444557) with the listed TaqMan probes. In all experiments, the TaqMan reference probe for TBP expression was used as endogenous loading control. To assess log2 changes in expression, −ΔΔCt values were determined for each condition compared to TBP reference.

### Immunoblotting (Western blots)

Total protein was harvested from cells or dissociated spheroids using RIPA lysis buffer (50 mM Tris HCl pH 7.5, 150 mM NaCl, 1% NP-40, 0.5% sodium deoxycholate, 0.1% SDS, cOmplete ULTRA protease inhibitors (Roche #5892791001). Briefly, dissociated cells were washed 3 times in cold PBS before resuspension in cold RIPA lysis buffer. After a 10 minute incubation on ice, tubes were briefly vortexed before sonication using a Bioruptor (5 cycles of 30 seconds ON/OFF on ‘high’ power). Samples were briefly vortexed before clearing by centrifugation at 18,000 x *g* for 15 minutes at 4 °C. Supernatant was transferred to a new pre-chilled tube. Protein yield was determined using the Pierce Detergent Compatible Bradford Assay Kit (Thermo Fisher #23246) according to manufacturer’s instructions. Lysates were separated on Novex NuPAGE Bis-Tris 4-12% polyacrylamide gels with MOPS SDS running buffer (Thermo Fisher #NP0366 and NP0001) and transferred to 0.45 μm PVDF membrane (Millipore #IPFL00010). Membranes were sectioned according to molecular weight range of target protein. After blocking (5% blotto or 5% BSA in TBS-T), blots were incubated in primary antibody overnight at 4 °C, washed 3 times in TBS-T, and probed with horseradish-peroxidase-labelled secondary antibody for 1-2 hours. Detection was performed using Biorad Clarity Western (#1705060) or Max (#1705062) ECL substrate on a Bio-Rad ChemiDoc MP imager. A list of antibodies and dilutions are available in the table.

### Immunohistochemistry and histology

Spheroids were cryopreserved and sectioned for staining as follows: after growth for 9 days, spheroids were fixed with 10% formalin for 30 minutes, washed 2X in PBS, and embedded in Tissue-Tek O.C.T. Compound for storage at −80 °C until sectioning. For KI-67 staining, 6 μm spheroid sections were mounted on positively charged slides. After heat-induced epitope retrieval, slides were incubated with KI-67 antibody and detection was performed using an HRP-polymer secondary antibody followed by 3,3’-Diaminobenzidine (DAB) staining.

### Spheroid fractionation

Spheroids were prepared as described above. After 9 days of growth, 75 μL of growth media was removed from each well and 50 μL of 4 μM CellTrace Far Red (in DMEM high glucose, no supplements) was added. Unstained, dissociated spheroids served as a negative control during FACS. Spheroids were incubated for 40 minutes under normal culture conditions (37 °C, 5% CO_2_). Any remaining dye not taken up by spheroids was neutralized with 100 μL culture media (+ serum) prior to spheroid dissociation. Spheroids were dissociated into a single-cell suspension by washing once with PBS and mechanically dissociating with TrypLE Express.

Cell suspensions were subsequently pelleted, resuspended in PBS + 1% FBS, and filtered through a 35 μm strainer (Falcon #352235) to remove any cell aggregates prior to FACS. After gating for live, single cells, FACS was performed using a BD FACS AriaIIu instrument at 630 nm excitation / 661 nm emission. Cells were sorted into two populations representing inner (dimmest 20% of population) and outer (brightest 20% of population). For RNA-seq, spheroid fractions were sorted directly collected into Sigma TRI Reagent LS, and RNA was harvested according to manufacturer’s instructions. For ATAC-seq, cells were collected into culture media and processed for ATAC-seq as described below.

### Visualization of spheroid staining method

Spheroids were cultured for 9 days and stained with CellTrace Far Red as described above. After staining, spheroids were embedded in Tissue-Tek O.C.T. compound for cryopreservation at −80 °C until sectioning. After cryosectioning on a CryoStar NX70 (Thermo Fisher), spheroid sections were mounted onto positively charged slides, counterstained with DAPI, and imaged using a Nikon ECLIPSE Ti2 inverted microscope.

### RNA sequencing and analysis

Total RNA obtained from TRIzol extraction was subjected to polyA enrichment, fragmentation, and random primer cDNA synthesis as previously performed^47^. Sequencing on all libraries was performed using Illumina sequencing on either a NextSeq 500 high output flow cell, or Illumina NovaSeq 6000 flow cell according to standard Illumina protocols.

RNA-seq reads obtained were processed by mapping to the hg38 reference human genome using HISAT2 2.1.0^82^ and reads with mapping quality <10 were discarded, leaving only high-quality reads. Reads within genes were counted using HTseq^83^ and these counts were processed using DESeq2 with default parameters^84^. Log2 fold changes were calculated using maximum a posteriori estimation using a zero-mean normal prior (Tikhonov-Ridge regularization). FDR-corrected P values were calculated using the Benjamini-Hochberg procedure.

### Identification of 3D-specific transcripts

All 2D, 3D samples and TCGA samples were separately pooled and analyzed using DESeq2 as described above. Inclusion criteria for the 3D-specific signature were as follows: An individual gene was considered as a member of this signature if altered transcription between 2D and TCGA samples was detected with fold change >1.5-fold in either direction and FDR-adjusted P<0.10 in at least one cell line. Furthermore, for each cell line, each gene was assigned a penalty if the expression fold change in 3D was opposite in magnitude as the expression fold change in primary tumors, compared to the respective 2D condition. We assigned this penalty to be the product of the fold changes in 3D and in primary tumors, which was negative in the case of discordant changes. For each gene, this penalty was summed across the 7 cell lines, and only genes with summed penalty values < 1 were considered. This strategy penalized discordant changes while permitting small numbers of low-magnitude mismatches, resulting in 1,912 consensus transcripts presented in Figure 1.

### Single-cell RNA sequencing (scRNA-seq)

3D spheroids were cultured for 9 days. Single-cell suspensions were obtained by washing once with PBS and mechanically dissociating with TrypLE Express. Cell suspensions for each cell line were combined to obtain a cell suspension of 1×10^6^ cells/mL in PBS + 0.04% BSA (Sigma #SRE0036). Cell suspensions were processed for single-cell RNA-seq libraries using the 10X Genomics Chromium platform according to the 3’ RNAseq V2 library kit instructions. Sequencing was performed on an Illumina NovaSeq 6000 sequencer according to standard protocols.

Sequenced scRNA-seq libraries were processed using the 10x Cell Ranger Pipeline 2.0.1 by mapping to the hg19 genome. The average mRNA expression level from each hallmark gene set in MSigDB cell was analyzed using the Single-Cell Signature Explorer^85^ to calculate an overall single-cell expression intensity score for each gene set.

### Inference of spatial positions of cells for scRNA-seq

Single-cell pseudospace trajectories were constructed as previously described^86,87^ based on differentially expressed genes in inner and outer fractions obtained using conventional RNA-seq of FACS-sorted samples. Briefly, differentially expressed genes between inner and outer spheroid fractions (|log2FC| >1 and adjusted P<0.05) from the FACS-based spheroid fractionation RNA sequencing (described above) were selected to build a pseudospace trajectory using Monocle2. The mRNA expression level of each individual single cell was first normalized by Seurat (3.1.1), then the dimensionality of the expression matrix was reduced using the “DDRTree” method. The two pseudospace trajectories of SMARCA4 and empty vector were aligned with dynamic time wrapping to a common pseudospace axes with package DTW (1.21-3). Single cells were then ranked separately along the pseudospace and aggregated into 100 bins, each corresponding to 1 %ile position bins. The average mRNA expression level from each bin was then analyzed using the Single-Cell Signature Explorer^85^ to calculate the signature score of all the KEGG and Hallmark gene sets. Heatmaps were plotted using the ComplexHeatmap (2.0.0) R package. Rank pseudospace coordinates were extrapolated into three dimensions and visualized using the rgl R package.

### Cell cycle phase scoring of scRNA-seq

The cell cycle phase of individual cells was inferred as described previously^88^. Briefly, cells were assigned a cell cycle phase score based on transcript- and phase-centered expression levels of cell cycle phase specific genes. Using these cell cycle phase signature scores, the cell cycle phase was assigned based on the highest cell cycle phase score.

### ATAC sequencing and analysis

ATAC libraries were prepared from separately cultured MCTS samples using the OmniATAC protocol^89^. Briefly, upon MCTS dissociation, cell nuclei were obtained by resuspending cells in 50 μl of lysis buffer (0.1% Tween-20, 0.1% NP-40 and 0.01% Digitonin in RSB buffer (10 mM Tris-HCl pH 7.5, 10 mM NaCl, 3 mM MgCl2)) and incubation on ice for 3 min. The lysis buffer was washed out by RSB buffer supplemented with 0.1% Tween-20. Subsequently, nuclei were pelleted by centrifugation for 10 min at 500g, resuspended in 50 μl of Transposition mix (2.5 μl Tagment DNA enzyme (Illumina), 25 μl of Tagmentation DNA buffer (Illumina), 0.1% Tween-20 and 0.01% Digitonin) and incubated for 30 min at 37 °C. DNA was purified with a MinElute PCR purification kit (Qiagen), and libraries were amplified by PCR with barcoded Nextera primers (Illumina) using 2X NEBNext High-Fidelity PCR Master Mix (NEB). For sequencing, libraries were size-selected with AMPure XP beads (Beckman Coulter) for fragments between ~100 and 1,000 bp in length according to the manufacturer’s instructions. Sequencing was performed using paired-end reads on an Illumina NextSeq 500 high output flow cell.

### Transcription factor motif profiling

Motif profiling was performed by identifying transcription factor motifs over-represented in SMARCA4-increased ATAC-seq peaks. Enrichment was measured using HOMER^90^ (version 4.11). Briefly, 424 transcription factor motifs within the HOMER database were examined for differential enrichment within SMARCA4-dependent peaks. To identify increased or unchanged peaks, differential peak calling was performed using DESeq2 using default parameters, and requiring fold changes >1.5-fold in either direction and FDR-adjusted P values <0.10. P values for enrichment were obtained by measuring the frequency of their motif presence in increased peaks compared to unchanged peaks using the hypergeometric test.

### Single-cell ATAC sequencing (scATAC-seq)

Cell nuclei were obtained following the same protocol as described above for ATAC-sequencing. Isolated nuclei were processed for single-cell ATAC-seq library using the 10X Genomics Chromium platform according to the manufacturer’s instructions. Sequencing was performed on an Illumina NovaSeq 6000 sequencer according to standard Illumina protocols.

scATAC-seq samples were processed with the Cell Ranger ATAC pipeline (version 1.1.0) with reference genome hg38. Cells were filtered based on following criteria: cells with <15% of reads in peaks, blacklist ratio>0.01, nucleosome, signal>10 and transcription start site enrichment score<2 were removed from further analysis by Signac (version 0.2.5).

### Inference of spatial positions of cells for scATAC-seq

Differentially accessible peaks between inner and outer fraction of spheroids identified by differential peak calls using DESeq2 from bulk ATAC-seq data were used to construct pseudospace trajectories in scATAC-seq with Monocle2 (version 2.12.0), using the same procedure as for scRNA-seq described above. Rank pseudospace coordinates were extrapolated into three dimensions and visualized using the rgl R package.

### Correlation of SMARCA4 targets with neighboring peaks

*Cis*-coaccessibility network (CCAN) boundaries were obtained from scATAC-seq data using Cicero^51^ (version 3.12) with default parameters. SMARCA4 target sites were defined as sites with SMARCA4-dependent accessibility changes from bulk ATAC-seq (fold change >1.5-fold in either direction and FDR-adjusted P<0.10), that also overlapped a detected peak from SMARCA4 ChIP-seq. Pairwise correlation of all peaks was measured within each CCAN boundary using Pearson correlation across 1%ile spatial bins. To assess the distribution of all positively or negatively correlated interactions, sites with root mean squared correlation values <0.2 across all pairwise interactions within the CCAN were discarded in order to remove globally invariant sites.

### Chromatin immunoprecipitation (ChIP) sequencing

ChIP libraries were prepared from separately cultured MCTS samples. MCTS were dissociated to single cell suspension and fixed for 10 min in 1% formaldehyde. Excess formaldehyde was quenched by the addition of glycine to 125 mM. Fixed cells were washed, pelleted, and snap-frozen using liquid nitrogen. Chromatin immunoprecipitation was performed as described previously^3^. Briefly, cell nuclei were isolated, resuspended in shearing buffer and sonicated in a Covaris E220 sonicator to generate DNA fragments between approximately 200 and 1,000 bp in length. Chromatin was then immunoprecipitated overnight at 4 °C with antibodies bound to Protein G Dynabeads. (Life Technologies). Chromatin was eluted and digestion and de-crosslinking was performed at 65 °C overnight, DNA was extracted with phenol-chloroform and precipitated with ethanol. Size selection was performed by extracting 200–400 bp DNA fragments before PCR amplification and DNA was extracted using MinElute cleanup kits (Qiagen). After PCR amplification, DNA quantified by Qubit fluorometric quantitation. Sequencing was performed using single-end reads on the Illumina NextSeq 500 (high output) instrument. Antibodies used in these studies include Anti-SMARCA4 (Cell Signaling, Cat# 49360S), ATF3 (Sigma-Aldrich) and NRF2 (Cell Signaling, Cat# HPA001562).

### Metabolite analysis

Spheroids cultured for 9 days were dissociated using TrypLE Express. Dissociated spheroids were collected as 5 independent replicates. Cell suspensions were washed three times with cold PBS and 5×10^6^ cells of each independent replicate were pelleted and stored at −80 °C. Pellets were processed for metabolite extraction by the BCM Metabolomics Core. Briefly, frozen cell pellets and quality control standards were thawed at 4 °C and lysed using >3 freeze/thaw cycles in liquid nitrogen and ice. To each individual extract, 750 μL of ice-cold methanol:water (4:1) and 20 μL spike-in internal standard was added followed by ice-cold chloroform and water in a 3:1 ratio. Cellular debris was removed by separating organic and aqueous layers independently and recombining. Samples were analyzed using a 6490 triple quadrupole mass spectrometer (Agilent Technologies, Santa Clara, CA) coupled to a HPLC system (Agilent Technologies, Santa Clara, CA) via single reaction monitoring (SRM) as described previously^91–96^.

#### Reagents and internal standards

High-performance liquid chromatography (HPLC)-grade ammonium acetate from Sigma, acetonitrile, methanol, chloroform, and water were procured from Burdick & Jackson (Morristown, NJ). Mass spectrometry-grade formic acid was purchased from Sigma-Aldrich (St Louis, MO). Metabolite standards and internal standards for assays include trans-zeatin (Sigma-Aldrich).

#### Separation of amino acid metabolites

ESI positive mode was used to measure the amino acids. For the targeted profiling (SRM), the RP chromatographic method employed a gradient containing water (solvent A) and acetonitrile (ACN, solvent B, with both solvents containing 0.1% Formic acid). Separation of metabolites was performed on a Zorbax Eclipse XDB-C18 column (50 × 4.6 mm i.d.; 1.8 μm, Agilent Technologies, CA) maintained at 37°C. The binary pump flow rate was 0.2 ml/min with a gradient spanning 2% B to 95% B over a 25 minute time period. Gradient: 0 min-2% B; 6 min-2% of B, 6.5 min-30 % B, 7 min-90% of B, 12 min-95% of B,13 min-2% of B followed by re-equilibration at end of the gradient.

#### Separation of glycolysis, TCA and pentose pathway metabolites

Tricarboxylic acid cycle (TCA) and glycolysis cycle metabolites were identified by using 5 mM ammonium acetate in water pH 9.9 as buffer A and 100% acetonitrile as buffer B using a Luna 3 μM NH2 100 Å chromatography column (Phenomenex, Torrance, CA). The gradient used: 0-20 min-80% B (0.2 ml/min); 20-20.10 min-80% to 2% B; 20.10-25 min-2% B (0.3 ml/min); 25-30 min-80% B (0.35 ml/min); 30-35 min-80% B (0.4 ml/min); 35-38 min-80% B (0.4 ml/min); followed by re-equilibration at the end of the gradient to the initial starting condition 80% B at 0.2 ml/min. All identified metabolites were normalized to a spiked internal standard.

### Glucose consumption assay

Glucose consumption was measured using the ACCU-CHECK Performa (Roche) glucose meter and glucose strips. For this assay, 2 μL of culture media was removed from spheroid culture and measured. This measurement was compared to initial media-only readings to obtain glucose consumed.

### Reactive oxygen species (ROS) measurements

ROS levels were measured in live, intact spheroids using the CellROX assay kit (Invitrogen) following the manufacturer’s instructions. Briefly, spheroids were formed and cultured as described above. On day 10, 2 μL of 75 μM CellROX Deep Red Reagent in DMSO was added directly to wells containing spheroids to obtain a final concentration of 1 μM CellROX. *tert*-butyl hydroperoxide (Sigma-Aldrich) served as a positive control for ROS. After incubation for 1 hour under normal culture conditions (37 °C, 5% CO2), plates were read on a Cytation5 fluorescent plate reader (far red = 640/20 excitation, 681/20 emission; GFP = 540/18 excitation, 579/20 emission), with ROS levels determined by the ratio of far red over GFP fluorescence intensity.

### Cystine uptake

To assess Cystine uptake, cells from day 9 spheroids were stained in cystine-free media supplemented with 1 uM Cystine-FITC (EMD Millipore catalog #SCT047) for 30 minutes and washed with PBS before resuspension in PBS 1% FBS. Cystine-FITC uptake was subsequently measured by flow cytometry using a BD CantoII analyzer at 488 nm excitation / 530 nm emission for each condition.

### Metabolite dropout assays

For glutamine/glucose/cystine dropout experiments, spheroids were prepared as described above, but were seeded in media with variable concentrations of glutamine, glucose, or cystine. In particular, media was prepared using DMEM, high glucose, no glutamine, no methionine, no cystine (Gibco) or DMEM no glucose, no glutamine (Gibco) supplemented with 10% dialyzed FBS (Sigma-Aldrich), 1 mM sodium pyruvate (Gibco), 10 mM HEPES (Gibco) 1X Pen/Strep (Gibco) and appropriate concentrations of methionine (Sigma-Aldrich), L-Glutamine (Thermo Fisher Scientific), Glucose (Thermo Fisher Scientific) and L-Cystine (Sigma-Aldrich). Spheroids were cultured for 9 days prior to assessment of intact spheroid viability.

### Measurement of reducing capacity

Reducing capacity was measured using alamarBlue HS reagent (Thermo Fisher), a resazurin-based solution that functions as a cell health indicator. At experimental endpoints, 15 μL (10% culture volume) of alamarBlue HS reagent was added directly to each well. Plates were incubated at 37 °C for 2-3 hours and read at 560/590 (Ex/Em) using the Cytation5 fluorescent plate reader. For drug screens, spheroids were cultured for 3 days prior to drug treatment for 72 hours when cell viability was measured using alamarBlue HS reagent.

### Assessment of apoptosis by flow cytometry

Spheroids were grown for 3 days prior to treatment with 0 μM or 400 μM tBHP and subsequently cultured for 72 h before harvesting for flow cytometry. Apoptosis was assessed in dissociated spheroids using Annexin V-FITC Apoptosis Detection Kit (Sigma-Aldrich) according the manufacturer’s instructions. Cell death was measured using a BD CantoII analyzer at 488 nm excitation / 530 nm for Annexin V-FITC, 561 nm excitation / 568 nm emission for PI and 635 nm excitation / 780 nm emission was used as a compensation control for autofluorescence.

### Measurement of lipid peroxidation

To assess lipid peroxidation, day 9 spheroids were dissociated and stained for 30 minutes with Image-iT Lipid Peroxidation Sensor C11-BODIPY (Thermo Fisher) at a final concentration of 10 μM according to the manufacturer’s instructions. Cells were subsequently washed three times with PBS and allowed to rest prior to measurement to permit accumulation of oxidized dye. To compare the extent of lipid peroxidation, fluorescence was measured at two separate wavelengths: excitation/emission of 584/625 nm for the reduced dye, and excitation/emission of 488/510 nm for oxidized dye using the Cytation5 fluorescent plate reader.

### Cell growth and viability measurement using Trypan blue

Spheroids were grown for 3 days prior to drug and/or tBHP treatment. To ensure that each concentration of drug received equivalent amounts of DMSO, the following procedure to dilute drugs was employed: For each applied concentration, drugs were diluted in DMSO to 1000x the final concentration, then further diluted 100-fold in culture media to yield a 10x stock for each intended concentration. This 10x stock was added to individual wells containing spheroids to yield the final 1x applied concentration. For erastin and SC-1 experiments, spheroids were challenged with 200 uM of tBHP. For apoptosis and cell death experiments involving rescue, cells were exposed to with 400 uM of tBHP to induce cell death. Cells were cultured for up to 3 days before measuring cell viability with Trypan blue. To assess cell viability, spheroids were harvested and dissociated with trypsin. Trypsinization was terminated by addition of culture medium, and cell number/viability were determined based on Trypan Blue dye exclusion using a BioRad TC20 cell counter.

### Analysis of public LUAD cancer datasets

LUAD RNA-seq data was obtained from the TCGA and processed using the TCGAbiolinks R package^97^. Samples were separated into normal or primary tumor based on sample type codes N and TP, respectively. For all samples, RNA-seq data was normalized using FPKM (fragments per kilobase million reads), using the canonical (i.e. longest length) transcript length for each gene. The matrix of FPKM values across all TCGA LUAD samples was made by placing each FPKM value for each transcript as a row, with each column corresponding to an individual tumor specimen. For comparison of individual transcripts with cell line RNA-seq data, we normalized transcripts in our RNA-seq data from cell lines using FPKM as above.

For analysis of the correlation of SMARCA4 expression with all other genes entirely within TCGA samples, RNA-seq counts for all transcripts were normalized using RSEM^98^ obtained using the TCGAbiolinks package.

### Analysis of TCGA expression profiles based on genetic alterations

For analysis of the influence of genetic alterations in TCGA data sets, only tumor samples that were analyzed for both mutation calls and copy number variation were considered. Individual genes were assigned by cBioPortal^99^ into “altered” or “unaltered” categories if samples had “deep” copy-number deletions (typically considered biallelic), high-functional impact missense mutations, truncating mutations, or chromosomal fusions.

Selectively enriched gene sets were obtained by manually confirming statistically significant enrichment of genes based on each category of genetic alteration. NRF2 genes were selected from the NFE2L2.V2 gene set. Genes associated with EGFR alterations were determined by identifying genes with significant enrichment in EGFR-altered tumors compared to EGFR-unaltered tumors. Inflammatory genes were selected from the HALLMARK_INTERFERON_GAMMA_RESPONSE gene set. For all genes and tumors, expression was log2 transformed and scaled to normal variance across rows and columns for presentation in the heat map.

### Statistical analysis

GraphPad Prism (5.0) or R (3.6.1) was used for statistical analyses. Gene expression data are presented as mean ±SEM. A two-sided Student’s T test was used for analyses of gene expression and comparisons of two groups. Drug response data are presented as mean ±SEM. Two-way ANOVA was used to compare drug response curves and spatial gene expression. Box and whiskers plots represent default settings: boxes span first and third quartiles, whiskers extend 1.5x inter-quartile ranges. Metabolomics data were log2-transformed and normalized with internal standards on a per-sample, per-method basis. Statistical comparison of metabolite abundance was performed using the two-sample T test in R. For all tests involving multiple comparisons, P-values were adjusted using the Benjamini-Hochberg false discovery rate (FDR) correction procedure.

### Data availability

All high-throughput sequencing data used in this project have been deposited in the Gene Expression Omnibus (GEO) database with SuperSeries accession number GSE146270.

### Code availability

Code for data analyses is available at https://github.com/hodgeslab/workflows.

## Acknowledgements

This study was supported by NIH grants T32ES027801 (E.A.S.), R00CA187565 (H.C.H.), R35GM137996 (H.C.H.), P30CA125123, S10OD016167, S10OD023469, S10OD025240, and P30EY002520. Support was also provided by CPRIT grants RR170036 (H.C.H.), RR160047 (O.V.), RP170002, RP170005, RP180672, RP120348, Gabrielle’s Angel Foundation (H.C.H.), the V Foundation (H.C.H.), intramural funds from the Dan L Duncan Comprehensive Cancer Center (DLDCCC), and Rice University Academy Fellowship (M.I.J.). We thank A. Jain, B. Ke, S. Nouraein, M. Sayeeduddin, P. Castro, and J. Sederstrom for technical services, as well as C. Walker for feedback.

## Author contributions

E.A.S. and H.C.H. conceptualized the study. K.C., E.A.S., M.L.C., M.I.J., O.V., and H.C.H. designed experiments. K.C., E.A.S., Y.S.C., M.L.C., C.C., Y.X., and M.I.J. collected data; E.A.S., Y.S.C., K.C., and M.I.J. analyzed data; N.P., L.M.S., R.C., R.T.R., and O.V. provided expert guidance and analytical services; K.C., E.A.S., and H.C.H. prepared figures and wrote the manuscript; all work was performed under the supervision of H.C.H.

## Competing interests

Authors declare no competing interests.

## Materials and Correspondence

Further information and requests for resources and reagents should be directed to the Corresponding Author, H. Courtney Hodges.

## Notes

### Competing Interest Statement

The authors have declared no competing interest.

