## Supplemental Information for "SMARCA4 regulates spatially restricted metabolic plasticity in 3D multicellular tissue"

**Supplemental text**

Summary of key resources

| **REAGENT or RESOURCE** | | **SOURCE** | | **IDENTIFIER** |
| --- | --- | --- | --- | --- |
| **Antibodies** | | | | |
| Mouse Anti-SMARCA4 (BRG1) (H-10)  (1:3000-1:5000) | | Santa Cruz Biotechnology | | Cat# sc-374197 |
| Rabbit Anti-ARID1A (D2A8U)  (1:1000) | | Cell Signaling Technology | | Cat# 12354S |
| Rabbit Anti-PBRM1  (1:2000) | | Bethyl Laboratories | | Cat# A301-591A |
| Rabbit Anti-BRD9  (1:2000-1:3000) | | Abcam | | Cat# ab137245 |
| Mouse Anti-β-Actin (GT5512)  (1:10000-1:20000) | | GeneTex | | Cat# GTX629630 |
| Rabbit Anti-SLC7A11/xCT (D2M7A)  (1:1000 in 5% BSA) | | Cell Signaling Technology | | Cat# 12691S |
| Mouse anti-rabbit IgG-HRP  (1:5000) | | Santa Cruz Biotechnology | | Cat# sc-2357 |
| Anti-mouse IgG-BP-HRP  (1:10000) | | Santa Cruz Biotechnology | | Cat# sc-516102 |
| TTF-1 / NKX2-1 Antibody (SPM150) | | NOVUS | | Cat# NBP2-34742PE |
| Brg1 (D1Q7F) Rabbit mAb | | Cell Signaling | | Cat# 49360 |
| NRF2 (D1Z9C) XP® Rabbit mAb | | Cell Signaling | | Cat# 12721 |
| Anti-ATF3 antibody produced in rabbit | | Sigma-Aldrich | | Cat# HPA001562 |
| **Chemicals, Peptides, and Recombinant Proteins** | | | | |
| Puromycin | | Fisher Scientific | | Cat# BP2956100 |
| Hygromycin B, 50 mg/mL | | EMD Millipore | | Cat# 400052 |
| Geneticin (G418 sulfate), 50 mg/mL | | Gibco | | Cat# 10131035 |
| cOmplete ULTRA | | Roche | | Cat# 5892791001 |
| Dimethyl sulfoxide (DMSO) | | Sigma | | Cat# D2650 |
| DMEM, high glucose | | Gibco | | Cat# 11960-051 |
| DMEM, high glucose, no glutamine, no methionine, no cystine | | Gibco | | Cat# 21013024 |
| DMEM no glucose, no glutamine | | Gibco | | Cat# 64319-25G-F |
| Advanced DMEM/F-12 | | Thermo Fisher | | Cat# 12634-010 |
| RPMI 1640 | | Corning | | Cat# 15-040-CV |
| L-Methionine | | Sigma-Aldrich | | Cat# 64319-25G-F |
| L-Glutamine | | Thermo Fisher Scientific | | Cat# 25-030-081 |
| Glucose Solution | | Thermo Fisher Scientific | | Cat# A2494001 |
| L-Cystine | | Sigma-Aldrich | | Cat# C8755-100G |
| Sodium Pyruvate (100 mM) (100X) | | Gibco | | Cat# 11360-070 |
| MEM Non-essential amino acids (NEAA)(100X) | | Gibco | | Cat# 11140-050 |
| GlutaMAX (100X) | | Gibco | | Cat# 35050-061 |
| HEPES (1 M) | | Gibco | | Cat# 15630-080 |
| Trypsin-EDTA (0.25%), phenol red | | Gibco | | Cat# 25200-056 |
| TrypLE Express (1X), phenol red | | Gibco | | Cat# 12605-010 |
| PBS-CMF, pH 7.4 | | Gibco | | Cat# 10010-23 |
| Matrigel Matrix, Growth Factor Reduced | | Corning | | Cat# 354230 |
| TRIzol Reagent | | Invitrogen | | Cat# 15596026 |
| TRI Reagent LS | | Sigma-Aldrich | | Cat# T3934 |
| Fetal Bovine Serum (FBS) | | Corning | | Cat# 35-010-CV |
| Fetal Bovine Serum (FBS), dialyzed | | Sigma-Aldrich | | Cat# F0392 |
| Trypan Blue Solution, 0.4% | | Gibco | | Cat# 15250061 |
| Agarose | | Thermo Fisher | | Cat# BP160-500 |
| Liproxstatin-1 | | Sigma-Aldrich | | Cat# SML1414 |
| Ferrostatin-1 | | Sigma-Aldrich | | Cat# SML0583 |
| Deferoxamine mesylate salt | | Sigma-Aldrich | | Cat# D9533 |
| Necrostatin-1 | | Sigma-Aldrich | | Catt# N9037 |
| Z-VAD-FMK | | R&D Systems | | Cat# FMK001 |
| Erastin | | Sigma-Aldrich | | Cat# E7781 |
| SC-1 | | Sigma-Aldrich | | Cat# SML0858 |
| **Critical Commercial Assays** | | | | |
| High-Capacity cDNA Synthesis Kit | | Applied Biosystems | | Cat# 4368814 |
| Tumor Dissociation Kit, human | | Miltenyi Biotec | | Cat# 130-095-929 |
| LIVE/DEAD™ Viability /Cytotoxicity Kit | | Thermo Fisher | | Cat# L3224 |
| Pierce Detergent Compatible Bradford Assay Kit | | Thermo Fisher | | Cat# #23246 |
| TaqMan Fast Advance Master Mix | | Applied Biosystems | | Cat #444455 |
| TDE1 (Tagment DNA Enzyme) | | Illumina | | Cat# 1502786 |
| TD Tagment DNA Buffer | | Illumina | | Cat# 15027866 |
| NEBNext High-Fidelity PCR Master Mix | | New England BioLabs | | Cat# M0541L |
| MinElute PCR purification kit | | Qiagen | | Cat# 28004 |
| AMPure XP beads | | Beckman Coulter | | Cat# A63880 |
| CellROX Assay Kit | | Invitrogen | | Cat# C10491 |
| AlamarBlue HS | | Invitrogen | | Cat# A50100 |
| CellTrace Far Red Cell Proliferation Kit | | Invitrogen | | Cat# C34564 |
| Biotracker Cystine-FITC | | Sigma-Aldrich | | Cat# SCT047 |
| Annexin V-FITC Apoptosis Detection Kit | | Sigma-Aldrich | | Cat #APOAF |
| **Deposited Data** | | | | |
| Overall SuperSeries | | This study | | GEO: GSE146270 |
| **Public Data** | | | | |
| H3K4me1 ChIP-seq | | ENCODE Project | | ENCFF848DUD |
| H3K27ac ChIP-seq | | ENCODE Project | | ENCFF403ARJ |
| **Experimental Models: Cell lines** | | | | |
| A549 | | ATCC | | Cat# CCL-185, RRID:CVCL_0023 |
| H1299 | | ATCC | | Cat# CRL-5803, RRID:CVCL_0060 |
| H2030 | | ATCC | | Cat# CRL-5914, RRID:CVCL_1517 |
| H1975 | | ATCC | | Cat# CRL-5908  RRID:CVCL_1511 |
| H23 | | ATCC | | Cat# CRL-5800  RRID:CVCL_1547 |
| H358 | | ATCC | | Cat# CRL-5807, RRID:CVCL_1559 |
| H441 | | ATCC | | Cat# HTB-174, RRID:CVCL_1561 |
| Lenti-X 293T | | Clontech | | Cat# 632180 |
| **Oligonucleotides** | | | | |
| Taqman assay for *SMARCA4* | | Thermo Fisher Scientific | | Cat# Hs00231324_m1 |
| Taqman assay for *TBP* | | Thermo Fisher Scientific | | Cat# Hs00427620_m1 |
| Taqman assay for *SLC7A11* | | Thermo Fisher Scientific | | Cat# Hs00921938_m1 |
| Taqman assay for *HK2* | | Thermo Fisher Scientific | | Cat# Hs00606086_m1 |
| **CRISPR sgRNA oligos** | | | | |
| gRNA Target | Forward sequence | | Reverse sequence | |
| Non-targeting ctrl (NTC) | caccgGGGAGGTGGCTTTAGGTTTT | | aaacAAAACCTAAAGCCACCTCCCc | |
| ARID1A | caccgAATACTCACAGGCAAGCTGG | | aaacCCAGCTTGCCTGTGAGTATTc | |
| BRD9 | caccgCTTGACGGACAGTACCGCAG | | aaacCTGCGGTACTGTCCGTCAAGc | |
| PBRM1 | caccgGAAACCACTTCATAATAGTC | | aaacGACTATTATGAAGTGGTTTCc | |
| **Recombinant DNA** | | | | |
| pBABE-hsBRG1 | | Addgene | | Addgene_1959 |
| pRRL-CAG-EV-IRES-PURO | | This study | | N/A |
| pRRL-CAG-hsSMARCA4-IRES-PURO | | This study | | N/A |
| CMV-SLC7A11-sh926R-FLAG-IRES-Hygro | | Addgene | | Addgene_118702 |
| LentiCRISPRv2-neo | | Addgene | | Addgene_98292 |
| **Software and algorithms** | | | | |
| Fiji | | https://imagej.net/Fiji | | N/A |
| R (3.6.1) | | https://www.r-project.org | | N/A |
| Nikon NIS-Elements AR  (5.11.01) | | https://www.nikon.com/products/microscope-solutions/lineup/img_soft/nis-elements/nis-elements_hc.htm | | N/A |
| FlowJo (10) | | FlowJo, LLC | | N/A |
| CellRanger | | https://www.10xgenomics.com | | N/A |
| Monocle2 (2.12.0) | | http://cole-trapnell-lab.github.io/monocle-release/ | | N/A |
| Cell Ranger ATAC (1.1.0) | | https://www.10xgenomics.com | |  |
| bedtools (version 2.28) | | https://bedtools.readthedocs.io/en/latest/content/history.html | | N/A |
| ComplexHeatmap (version 2.5.3) | | https://jokergoo.github.io/ComplexHeatmap-reference/book/ | | N/A |

**Supplemental figures**

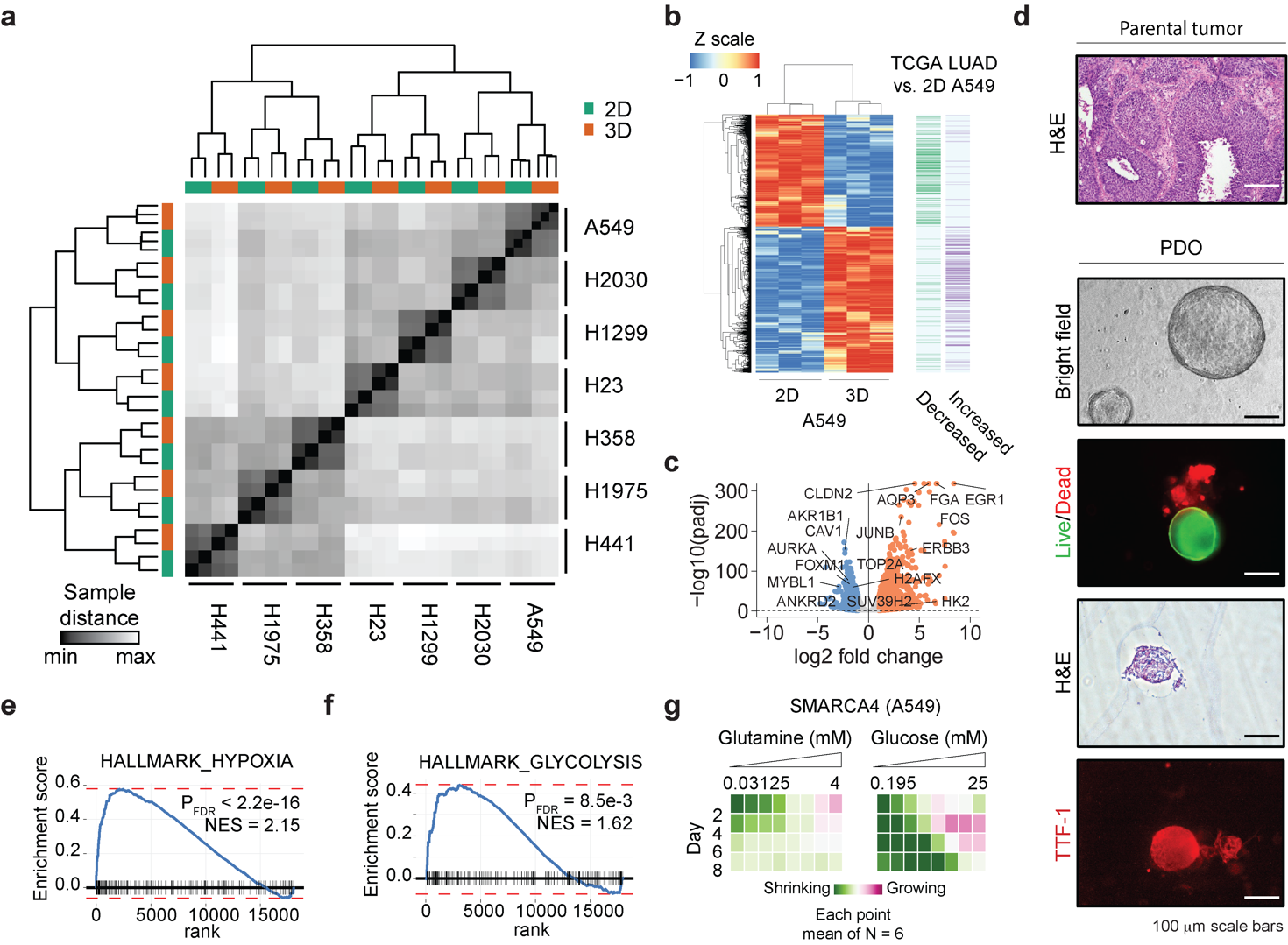

**Figure S1. Comparison of expression patterns of 2D and 3D lung adenocarcinoma (LUAD) cells.**

**(a)** Unbiased clustering of 2D and 3D LUAD samples based on their transcriptomes.

**(b)** Heatmap summary of gene expression changes determined by RNA-seq for A549 cells in 3D vs. 2D compared to differential expression of TCGA LUAD transcriptomes vs. 2D A549 cells.

**(c)** Volcano plot of gene expression changes for A549 cells in 3D vs. 2D from RNA-seq.

**(d)** Characterization of patient-derived organoid (PDO) derived from LUAD tumor.

**(e)** GSEA plot of 3D vs 2D A549 enrichment of hypoxia hallmark.

**(f)** GSEA plot of 3D vs 2D A549 enrichment of glycolysis hallmark.

**(g)** Change in growth of A549 spheroids expressing SMARCA4 over time in varying concentrations of glutamine and glucose.

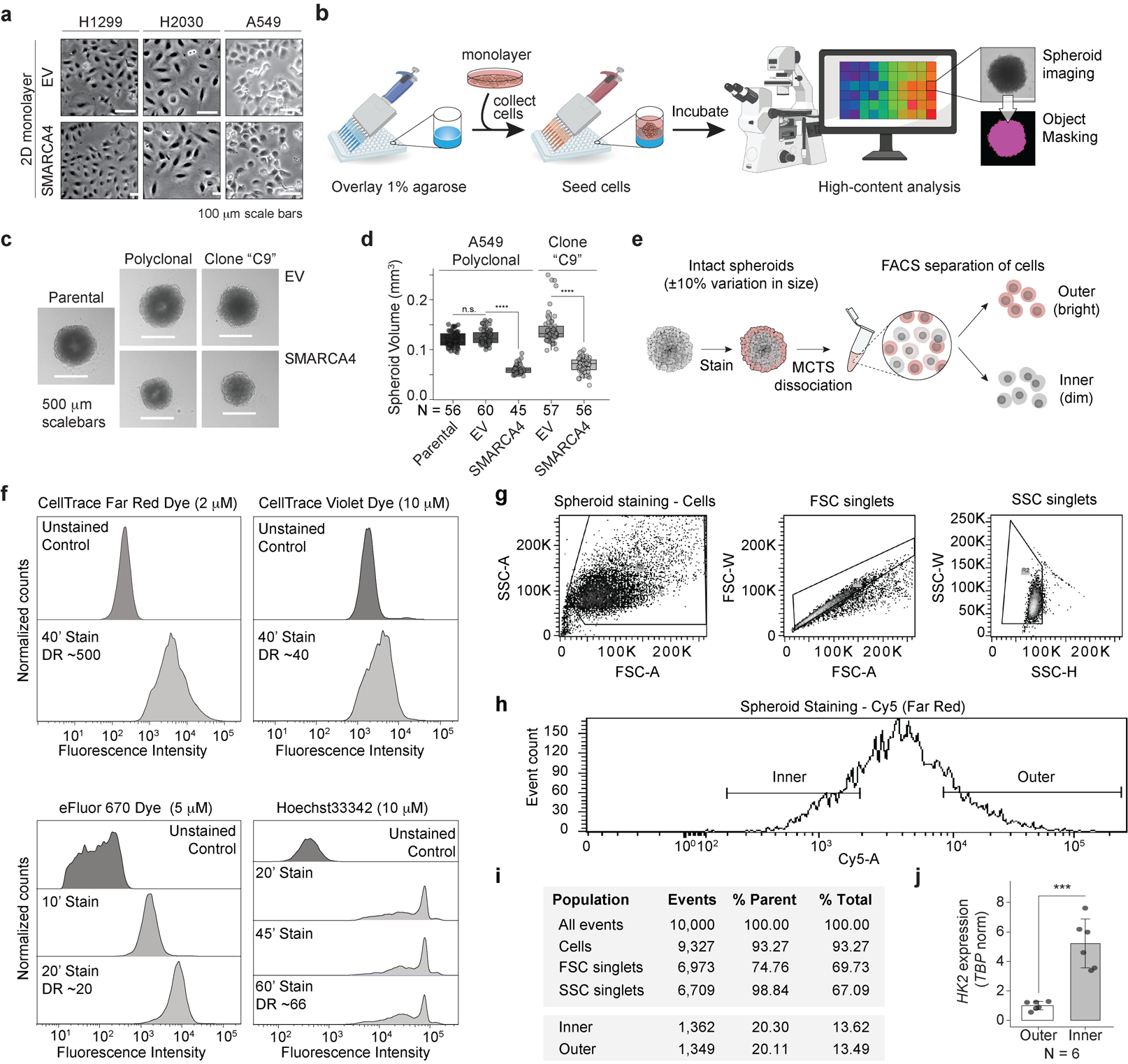

**Figure S2. Development of FACS-based approach for spatial reconstruction on MCTSs**

**(a)** Brightfield images of 2D culture of EV- or SMARCA4-expressing LUAD cell lines.

**(b)** Expanded workflow of high-content analysis of 3D spheroids.

**(c)** Brightfield spheroid images derived from parental A549 cells, or polyclonal and monoclonal A549 cells expressing EV or SMARCA4.

**(d)** Volumes of spheroids derived from parental A549 cells as well as EV- and SMARCA4-expressing A549 cells in polyclonal and monoclonal background (“C9”), seeded with equal cell numbers. **** P < 1e-4

**(e)** Workflow for spheroid staining and FACS separation of outer and inner spheroid fractions.

**(f)** Screening of spheroid staining methods by flow cytometry. CellTrace Far Red and Violet dyes, eFluor670 and Hoechst33342 staining of intact spheroids at different time points. CellTrace Far Red was selected for all studies for its high staining intensity and large dynamic range (DR).

**(g)** Gating strategy for forward (FSC) and side (SSC) scatter used to identify live singlet cells.

**(h)** Gating strategy for fluorescence activated cell sorting to separate cells from inner and outer spheroid fraction.

**(i)** Summary of cell abundances in each gate.

**(j)** *HK2* qPCR of staining/fractionation experiments. Mean ±SD plotted (N=6). ***P < 0.001

For all panels: EV, Empty vector

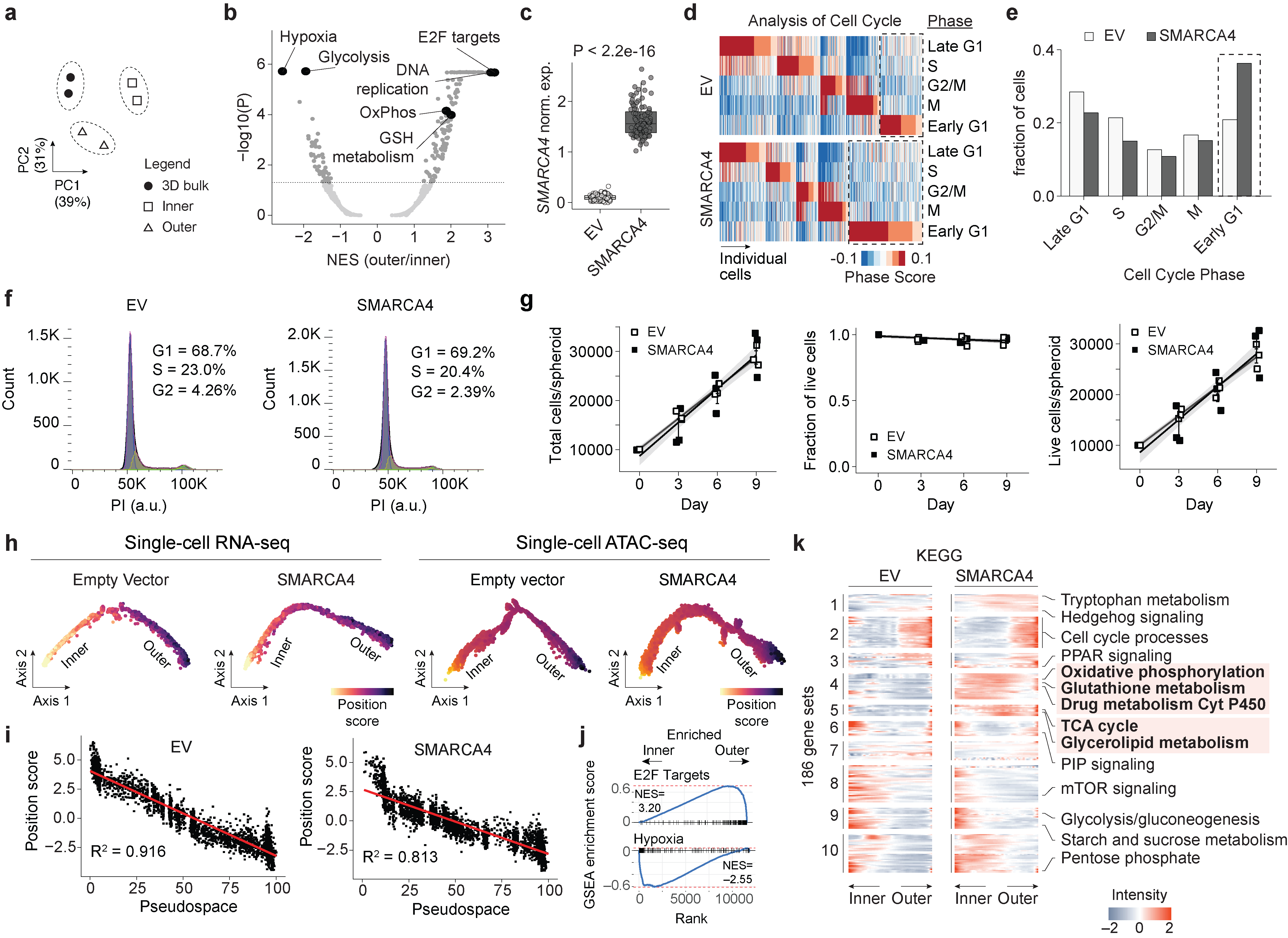

**Figure S3. Transcriptomic characteristics of sorted spheroids and single-cell RNA-seq.**

**(a)** Principal component analysis of bulk A549 spheroids RNA-seq compared to inner and outer sorted spheroid fractions.

**(b)** Volcano plot of Hallmark, KEGG, and Reactome GSEA of RNA-seq of outer versus inner sorted fractions of A549 spheroids.

**(c)** Normalized counts of *SMARCA4* expression across aggregated bins of single-cell RNA-seq.

**(d)** Cell cycle phase scoring and clustering of single-cell RNA-seq according to expression of cell cycle markers.

**(e)** Bar plot representation of cell fractions in each cell cycle phase calculated in (d).

**(f)** Cell cycle analysis of A549 cells in 3D spheroid culture using propidium iodide staining analyzed by flow cytometry.

**(g)** Cell growth of A549 cells in 3D spheroid culture measured by Trypan blue.

**(h)** Pseudospace trajectories of single cell RNA-seq and single-cell ATAC-seq for A549 spheroids derived from inner and outer spheroid fraction gene signatures.

**(i)** Pseudospace position of individual cells from single-cell RNA-seq is strongly correlated with correlation-based position score based on flow-sorted RNA-seq.

**(j)** GSEA of selected enriched gene sets comparing inner and outer fraction of A549 spheroids.

**(k)** Spatially resolved KEGG gene set enrichment for A549 spheroids. Identities of all gene sets are provided in **Supplemental Dataset 7**.

For all panels: EV, Empty vector

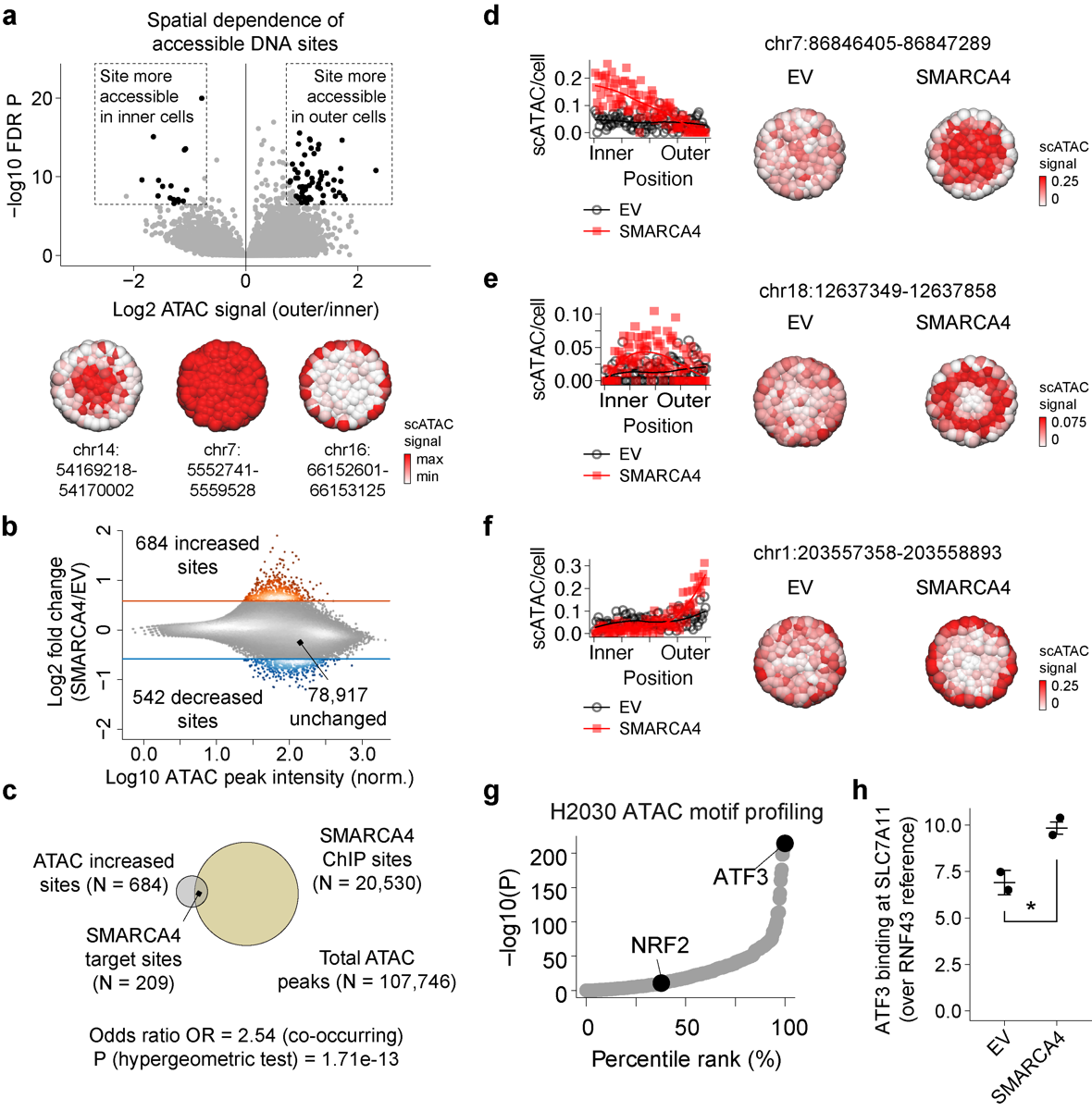

**Figure S4. SMARCA4-dependent regulation of accessibility in 3D LUAD spheroids**

**(a)** Volcano plot of sites with differential accessibility in outer versus inner spheroid fractions determined by ATAC-seq. Inferred spatially regulated accessibility patterns presented at bottom for example sites.

**(b)** Fold change of ATAC read density at individual sites across the genome comparing SMARCA4 vs. EV bulk ATAC-seq.

**(c)** Venn diagram identifying SMARCA4-bound target sites.

**(d-f)** Example of SMARCA4 target sites regulated in the inner (d) intermediate (e) or outer (f) portion of the spheroid. Accessibility patterns within spheroids are plotted and inferred spatial distribution presented on spheroid cartoons.

**(g)** Transcription factor motif accessibility enrichment in SMARCA4-increased ATAC sites measured using H2030 spheroids, ranked by P value.

**(h)** ChIP-qPCR of ATF3 binding at *SLC7A11* locus. * P < 0.05

For all panels: EV, Empty vector

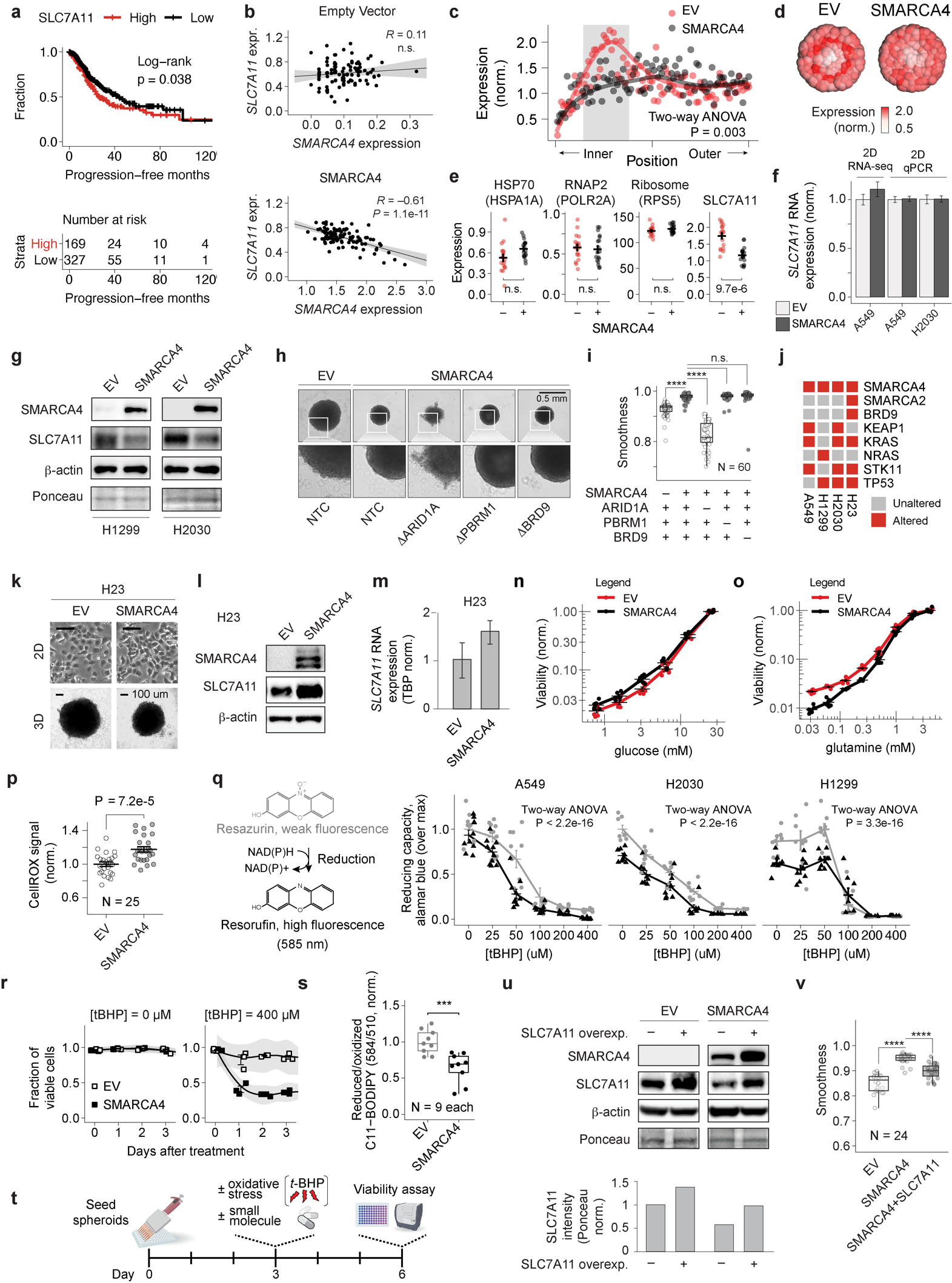

**Figure S5. SMARCA4 regulates SLC7A11.**

**(a)** Progression-free survival of LUAD patients based on SLC7A11 expression.

**(b)** Correlation between *SLC7A11* and *SMARCA4* expression in single cell RNA-seq.

**(c)** Spatial regulation of *SLC7A11* expression in A549 spheroids expressing EV or SMARCA4.

**(d)** Inferred spatial distribution of *SLC7A11* expression from panel (c).

**(e)** Expression of reference genes and *SLC7A11* levels within highlighted bins of panel (c).

**(f)** qPCR and RNA-seq of *SLC7A11* regulation in 2D A549 and 2D H2030 cells expressing EV or SMARCA4.

**(g)** Western blot of SLC7A11 expression in H1299 and H2030 3D spheroids.

**(h)** Brightfield images of indicated complex-specific knockouts in A549 spheroids. Inset zoom for comparison of edge smoothness spheroids. NTC, non-targeting CRISPR control.

**(i)** Spheroid edge smoothness measurements of BAF/PBAF/GBAF complex-specific knockouts in A549 spheroids expressing EV or SMARCA4. **** P < 0.0001

**(j)** Mutational status of select genes for LUAD cell lines used in this study.

**(k)** Brightfield images of H23 cells expressing EV or SMARCA4 in 2D and 3D cell culture.

**(l)** Western blot of SLC7A11 expression in H23 3D spheroids expressing EV or SMARCA4.

**(m)** qPCR of *SLC7A11* expression in 3D cultured H23 cells.

**(n)** Relationship of cell viability at day 9 and decreasing glucose dropout for A549 spheroids.

**(o)** Relationship of cell viability at day 9 and decreasing glutamine dropout for A549 spheroids.

**(p)** Levels of reactive oxygen species (ROS) in A549 spheroids by Cell ROX Deep Red staining. GFP fluorescence channel was used as an autofluorescence control.

**(q)** Measurement of resazurin reducing capacity with increasing concentration of tBHP.

**(r)** Cell viability of 3D A549 spheroids expressing EV or SMARCA4 treated with 0 or 400 μM *tert*-butyl hydroperoxide (tBHP) measured by Trypan blue.

**(s)** Lipid peroxidation measurements assessed by BODIPY-C11.

**(t)** Timeline for small molecule treatments in 3D spheroids

**(u)** Western blot of SLC7A11 protein levels upon ectopic SLC7A11 expression in EV- and SMARCA4-expressing A549. Densitometry of SLC7A11 normalized based on Ponceau staining.

**(v)** Comparison of spheroid edge smoothness measurements in A549 spheroids. **** P < 1e-4

For all panels: EV, Empty vector

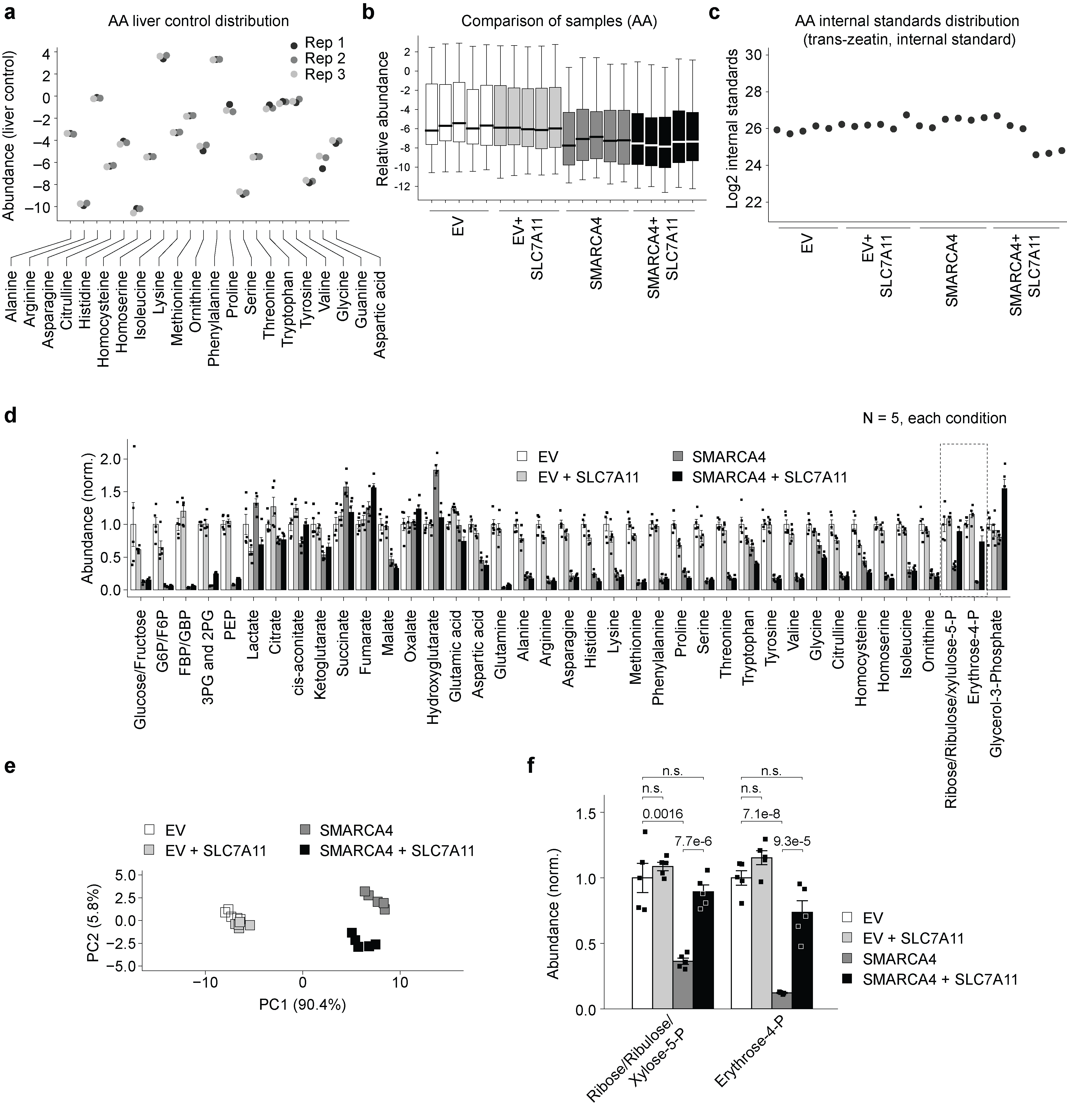

**Figure S6. Metabolomics and corresponding quality control data.**

**(a)** Reproducibility plot of amino acid measurements in standard liver control.

**(b)** Comparison of overall amino acid abundance across experimental conditions.

**(c)** Internal standard quality control metrics demonstrating consistent internal standard representation.

**(d)** Steady-state metabolites in A549 spheroids expressing EV or SMARCA4 ± ectopic expression of SLC7A11. N=5, each metabolite. SMARCA4 and EV data (without ectopic SLC7A11 expression) shown here are the same data as Figure 2p, reproduced here for clarity of comparison.

**(e)** Principal component analysis of steady-state metabolomics.

**(f)** Steady-state metabolites rescued by high levels of SLC7A11 in SMARCA4-expressing 3D spheroids (inset of panel (d) representing dotted region).

**Supplemental Datasets**

**Supplemental Dataset 1**. Identities and 3D-induced expression changes of 1,912 genes in 3D-specific LUAD expression signature. Tab-delimited text file.

**Supplemental Dataset 2.** Gene set enrichment analysis of 3D-induced expression changes in A549, H1299, and H2030 cells. Excel spreadsheet.

**Supplemental Dataset 3.** RNA-seq differential expression in inner, outer, and bulk 3D spheroids. Excel spreadsheet.

**Supplemental Dataset 4.** Inferred genome-wide spatial expression patterns in empty vector 3D A549 spheroids. Tab-delimited text file.

**Supplemental Dataset 5.** Inferred genome-wide spatial expression patterns in SMARCA4 3D A549 spheroids. Tab-delimited text file.

**Supplemental Dataset 6.** RNA-seq differential expression of bulk 3D spheroids expressing SMARCA4 vs. empty vector. Tab-delimited text file.

**Supplemental Dataset 7.** Identities of HALLMARK and KEGG gene sets examined for spatial expression changes. Excel spreadsheet.
